# Detection of low-frequency mutations and removal of heat-induced artifactual mutations using Duplex Sequencing

**DOI:** 10.1101/474353

**Authors:** Eun Hyun Ahn, Seung Hyuk Lee

## Abstract

We present a genome-wide comparative and comprehensive analysis of three different sequencing methods (conventional next generation sequencing (NGS), tag-based single strand sequencing (eg. SSCS), and Duplex Sequencing for investigating mitochondrial mutations in human breast epithelial cells. Duplex Sequencing produces a single strand consensus sequence (SSCS) and a duplex consensus sequence (DCS) analysis, respectively. Our study validates that although high-frequency mutations are detectable by all the three sequencing methods with the similar accuracy and reproducibility, rare (low-frequency) mutations are not accurately detectable by NGS and SSCS. Even with conservative bioinformatical modification to overcome the high error rate of NGS, the NGS frequency of rare mutations is 7.0×10^−4^. The frequency is reduced to 1.3×10^−4^ with SSCS and is further reduced to 1.0×10^−5^ using DCS. Rare mutation context spectra obtained from NGS significantly vary across independent experiments, and it is not possible to identify a dominant mutation context. In contrast, rare mutation context spectra are consistently similar in all independent DCS experiments. We have systematically identified heat-induced artifactual mutations and corrected the artifacts using Duplex Sequencing. All of these artifacts are stochastically occurring rare mutations. C>A/G>T, a signature of oxidative damage, is the most increased (170-fold) heat-induced artifactual mutation type. Our results strongly support the claim that Duplex Sequencing accurately detects low-frequency mutations and identifies and corrects artifactual mutations introduced by heating during DNA preparation.

## 1. Introduction

Next-generation sequencing (NGS) has rapidly transformed entire areas of basic research and therapeutic applications by making large scale genomic studies feasible through reduced cost and faster turnaround time [1,2]. NGS has been extensively used to study clonal (high-frequency) mutations, but not subclonal (low-frequency) mutations. A major impediment in investigating subclonal (low-frequency) mutations is that conventional NGS methods have high error rates (10^−2^ to 10^−3^), which obscure true mutations that occur less frequently than errors [3,4]. These subclonal mutations may account for the genetic heterogeneity of tumors and tumor recurrence, as well as provide a reservoir for the rapid development of resistance to chemotherapy [5].

Conventional sequencing technologies sequence only a single strand of DNA. In contrast, Duplex Sequencing examines both strands of DNA and scores mutations only if they are present on both strands of the same DNA molecule as complementary substitutions. This significantly reduces sequencing error rates to < 5 × 10^−8^ [6–9]. In the first report of Duplex Sequencing, accuracy and sensitivity of mutation detection were demonstrated mainly in M13mp2 bacteriophage by comparing untreated/control DNA and DNA incubated with hydrogen peroxide, a radical generator, in the presence of iron [6].

While overall frequencies and types of mutations from NGS, SSCS, and Duplex Sequencing have been compared in previous studies [5,10], those studies focused on detection limits of low-frequency mutations only and did not compare the sequencing methods’ ability to detect high-frequency mutations. In addition, influences of neighboring nucleotide base context on mutations (mutation context spectra) have not been investigated.

In the current study, we systematically compared mutation frequencies, types, positions, and sequence context spectra of the whole mitochondrial (mt) DNA in human breast epithelial cells using three different sequencing protocols: conventional NGS, tag-based single strand consensus sequencing (eg. SSCS), and Duplex Sequencing. We applied the three sequencing methods to categorize and analyze high-frequency and low-frequency mutations, separately. Furthermore, analyses were done with multiple independent DNA library preparation experiments of an identical biological sample to evaluate the detection consistency, reproducibility, and validity of each sequencing method. Heating samples, a common practice in preparing DNA for molecular biology experiments, can introduce such artifactual mutations [11]. Herein, we present heat-induced artifactual mutations identified using Duplex Sequencing and specific nucleotide contexts that contribute to a high level of heat-induced artifactual mutations.

## 2. Results

Duplex Sequencing generates both SSCS and DCS analysis results. In Duplex Sequencing, both strands of DNA are individually tagged and strands with identical tag sequence, the product of the same DNA template, are grouped together after PCR amplification. SSCS analysis differs from DCS analysis in that complementary tag sequences are not identified, and so complementary strands are not grouped [6]. The SSCS method represents a tag-based single strand sequencing procedure and is comparable to that of Safe Sequencing System (SafeSeqS) in that each single-stranded DNA molecule is uniquely labeled before PCR amplification, allowing strands of the same derivatives to be grouped [9,12].

The average number of nucleotides sequenced at each genome position (depth) of all NGS, SSCS, and DCS analyses were calculated as the total number of nucleotides sequenced divided by the mtDNA size of 16569 bases. The depths for NGS, SSCS, and DCS analyses for normal human breast cells and immortalized cells are presented in Table S1 and Table S2. The highest depths of NGS, SSCS and DCS that were processed under the same data processing conditions were 458441, 40421 and 6803, respectively (Table S1).

As an attempt to overcome the high error rates of NGS, more conservative bioinformatical conditions than those applied for SSCS and DCS, referred to as NGS (Q30^r^) hereinafter (See section: *Materials and Methods 4.3.2*), were applied to NGS datasets. Results of NGS (the same bioinformatical conditions as to SSCS and DCS) and NGS (Q30^r^) are presented in *Supplementary Figures S1 to S4*. Figure 1 through Figure 4 compare the results of NGS (Q30^r^) with those of SSCS and DCS.

**Figure 1.**
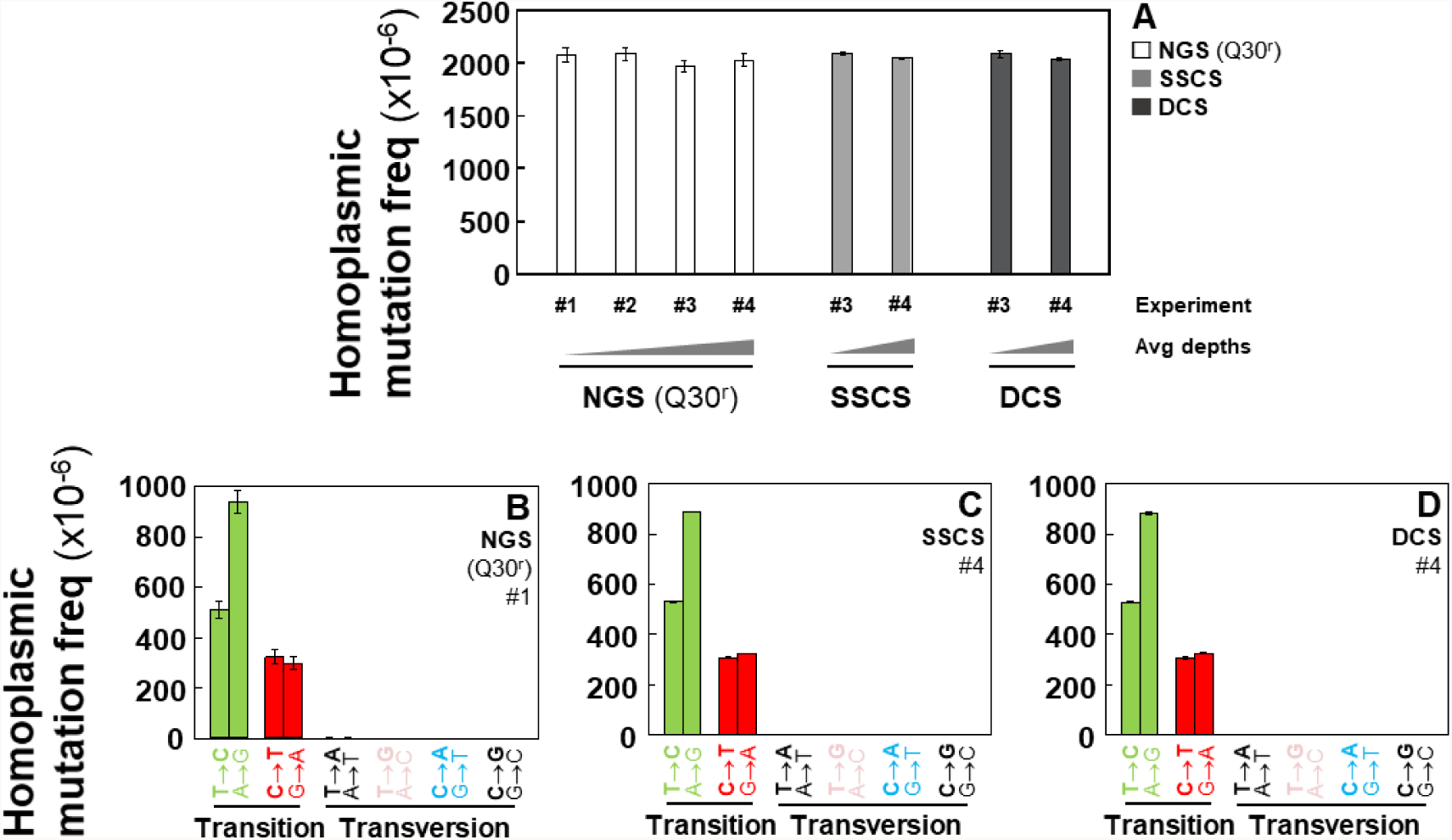
Frequencies of homoplasmic point mutations in the whole mtDNA of human breast immortalized cells. (**A**) Overall frequencies determined by performing NGS, SSCS, and DCS analyses. Homoplasmic mutation frequency of specific mutation types determined using (**B**) NGS, (**C**) SSCS, and (**D**) DCS analyses. Error bars represent the Wilson score 95% confidence intervals.

**Figure 2.**
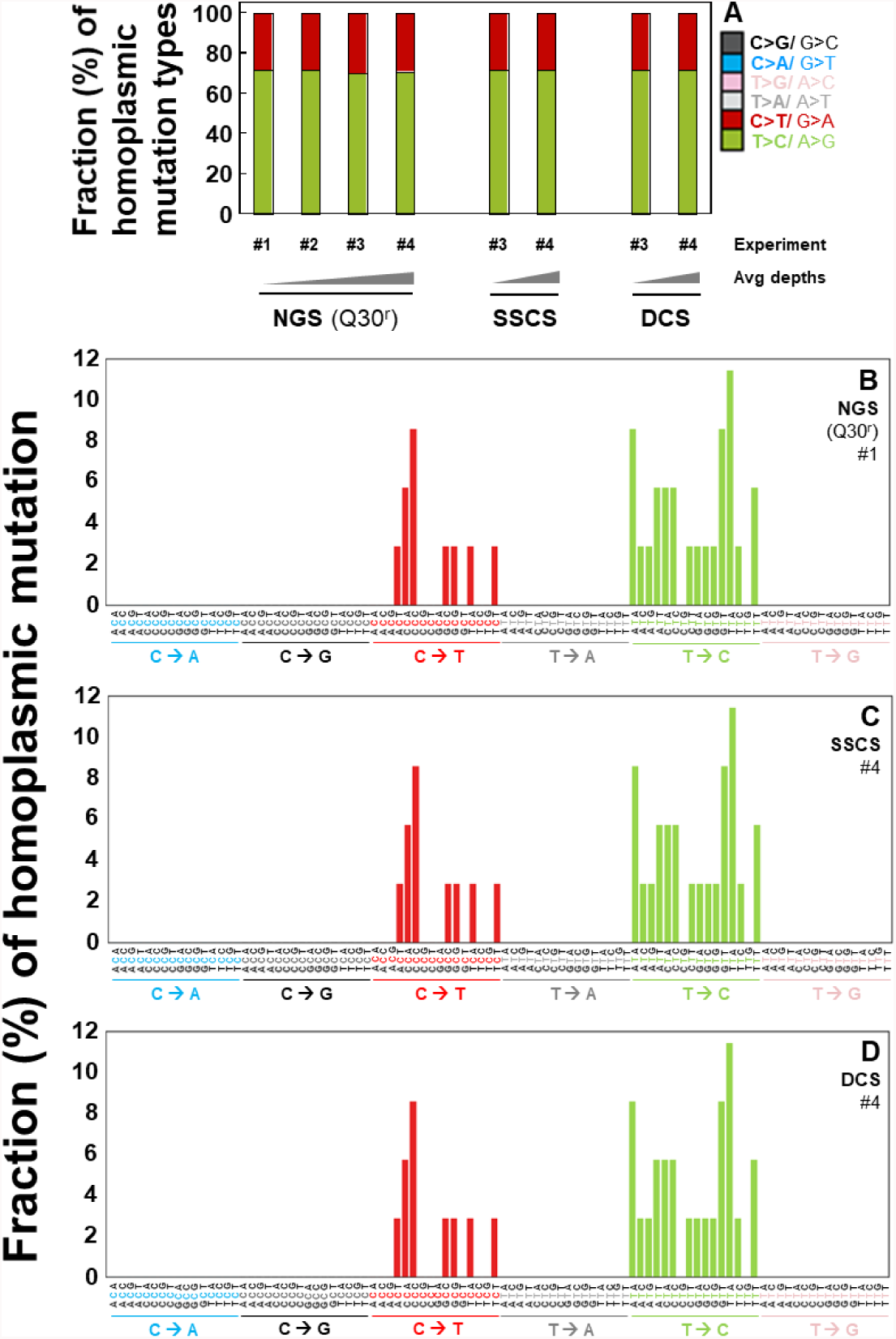
Fractions (%) of homoplasmic point mutation types and context spectra in the whole mtDNA of human breast immortalized cells. (**A**) Relative percentages of each mutation type in each experiment as determined by NGS, SSCS, and DCS analyses. Fractions of homoplasmic mutation context spectra determined by (**B**) NGS, (**C**) SSCS, and (**D**) DCS analyses.

**Figure 3.**
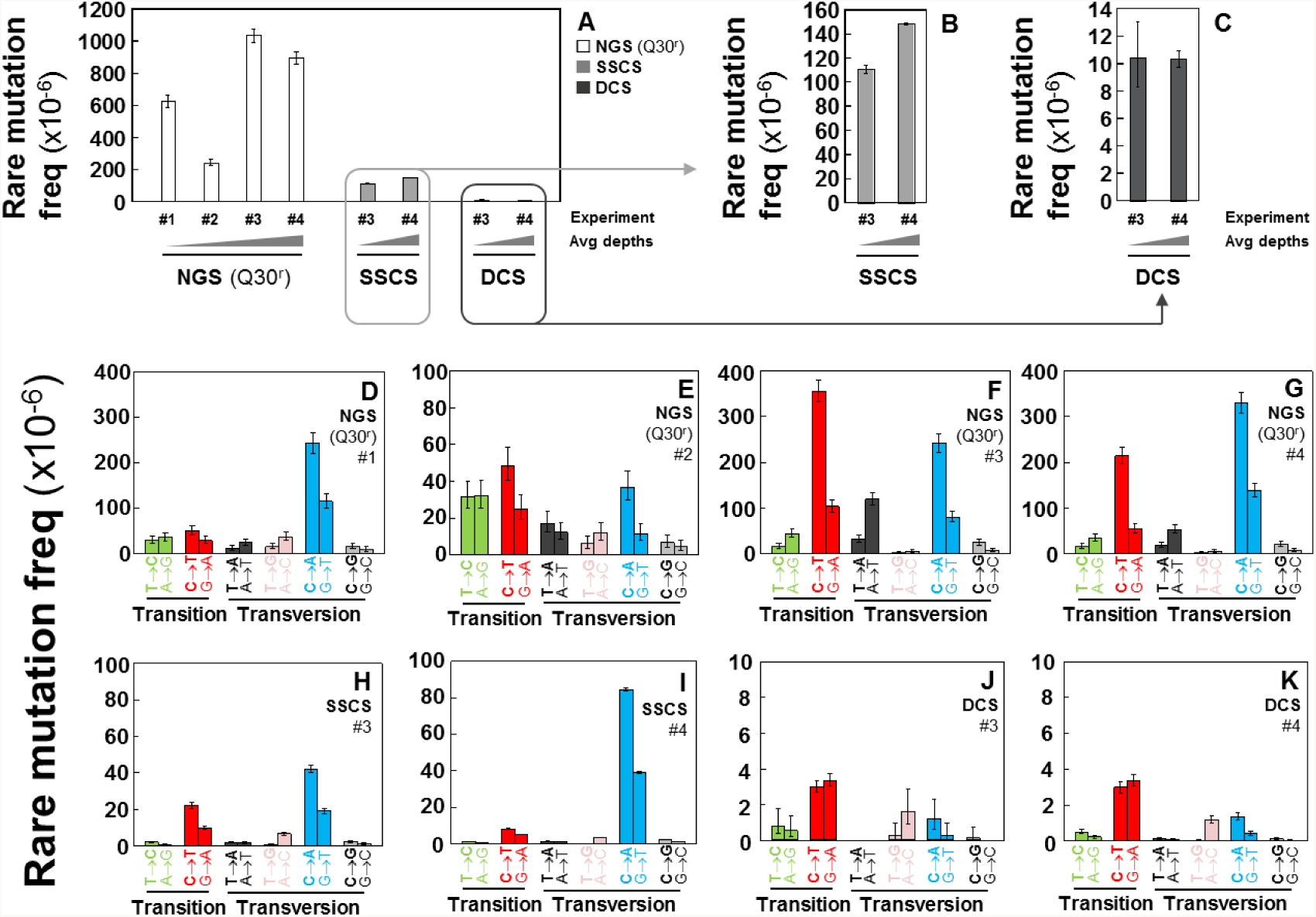
Frequencies of rare point mutations in the whole mtDNA of human breast immortalized cells. (**A-C**) Overall frequencies determined by performing NGS, SSCS, and DCS analyses. Rare mutation frequency of each mutation type as determined using (**D-G**) NGS, (**H,I**) SSCS, and (**J,K**) DCS analyses. Error bars represent the Wilson score 95% confidence intervals.

**Figure 4.**
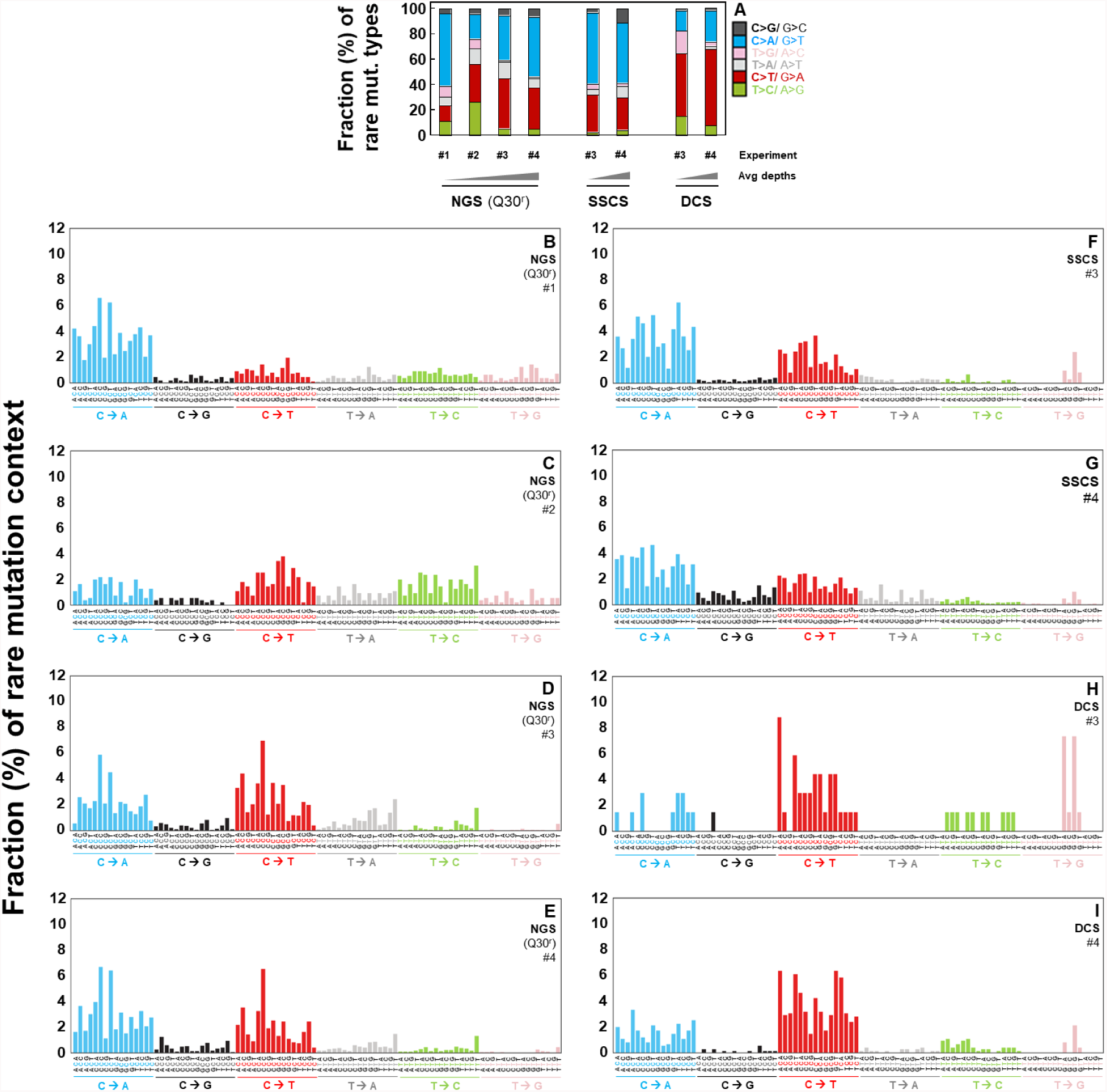
Fractions (%) of rare mutation types and context spectra in the whole mtDNA of human breast immortalized cells. **(A)** Relative percentages (%) of mutation types in each experiment. Fractions of rare mutation context spectra determined by performing (**B-E**) NGS, (**F,G**) SSCS, and (**H,I**) DCS analyses.

For this study, we have defined homoplasmic (90-100%: Figures 1,2,S1,S2, Table S3) and rare (0-1%: Figures 3-6, S3-S6, Table S4,S7) mutations based on the mutation occurrence (%) at each genome position. Mutation frequencies were calculated by dividing the number of variants by the total number of nucleotides sequenced.

**Figure 5.**
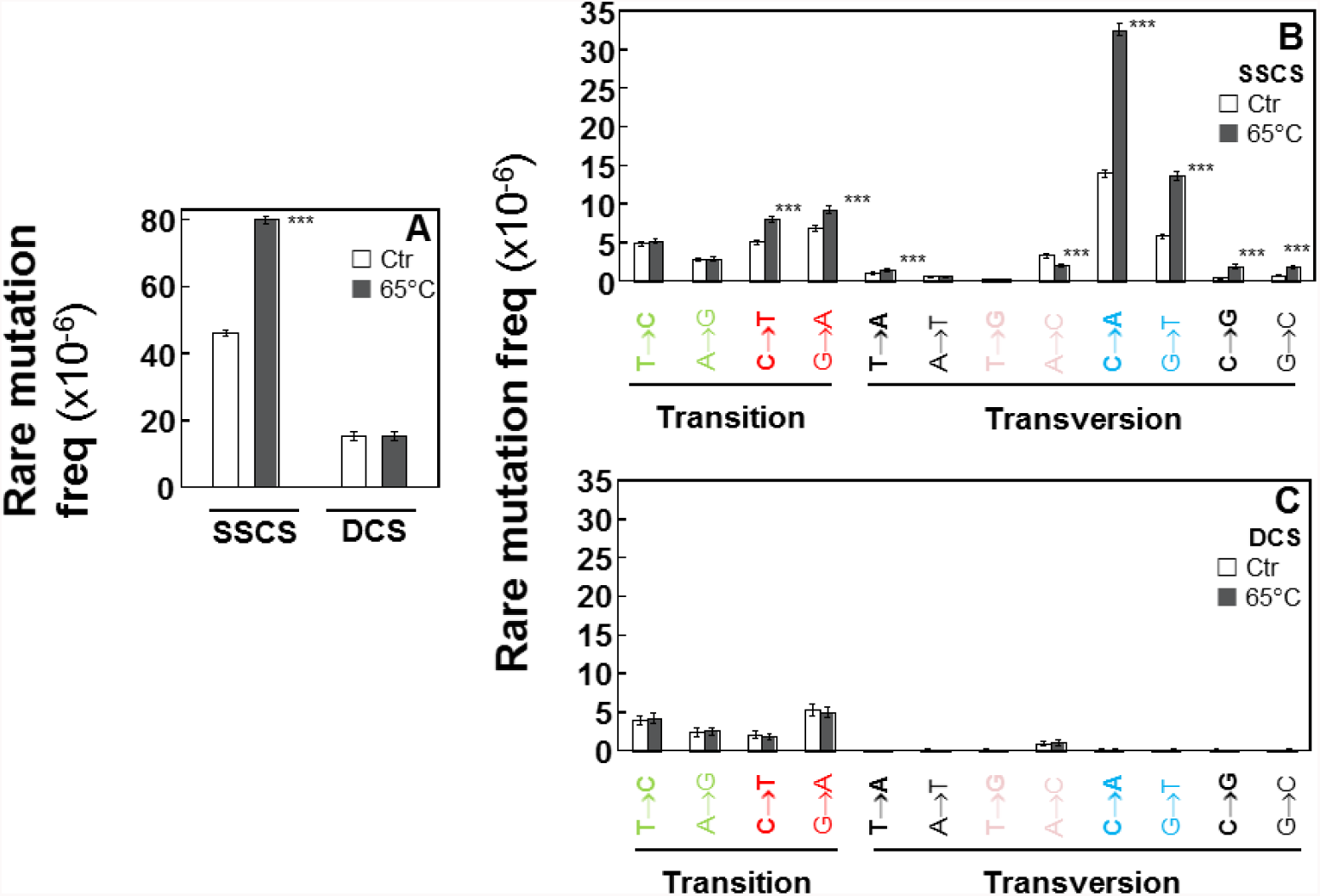
Frequencies of the heat-induced (65°C) artifactual mutations in human breast normal cells. Overall rare mutation frequency for heated versus control DNA as determined by SSCS and DCS analyses. **(B,C)** Frequencies of specific rare mutation types for heated versus control DNA. Error bars represent the Wilson score 95% confidence intervals. The significant differences in rare mutation frequencies between the control DNA and the heated DNA are indicated (\*\*\**p* < 5 × 10^−5^ by the Chi-square test).

**Figure 6.**
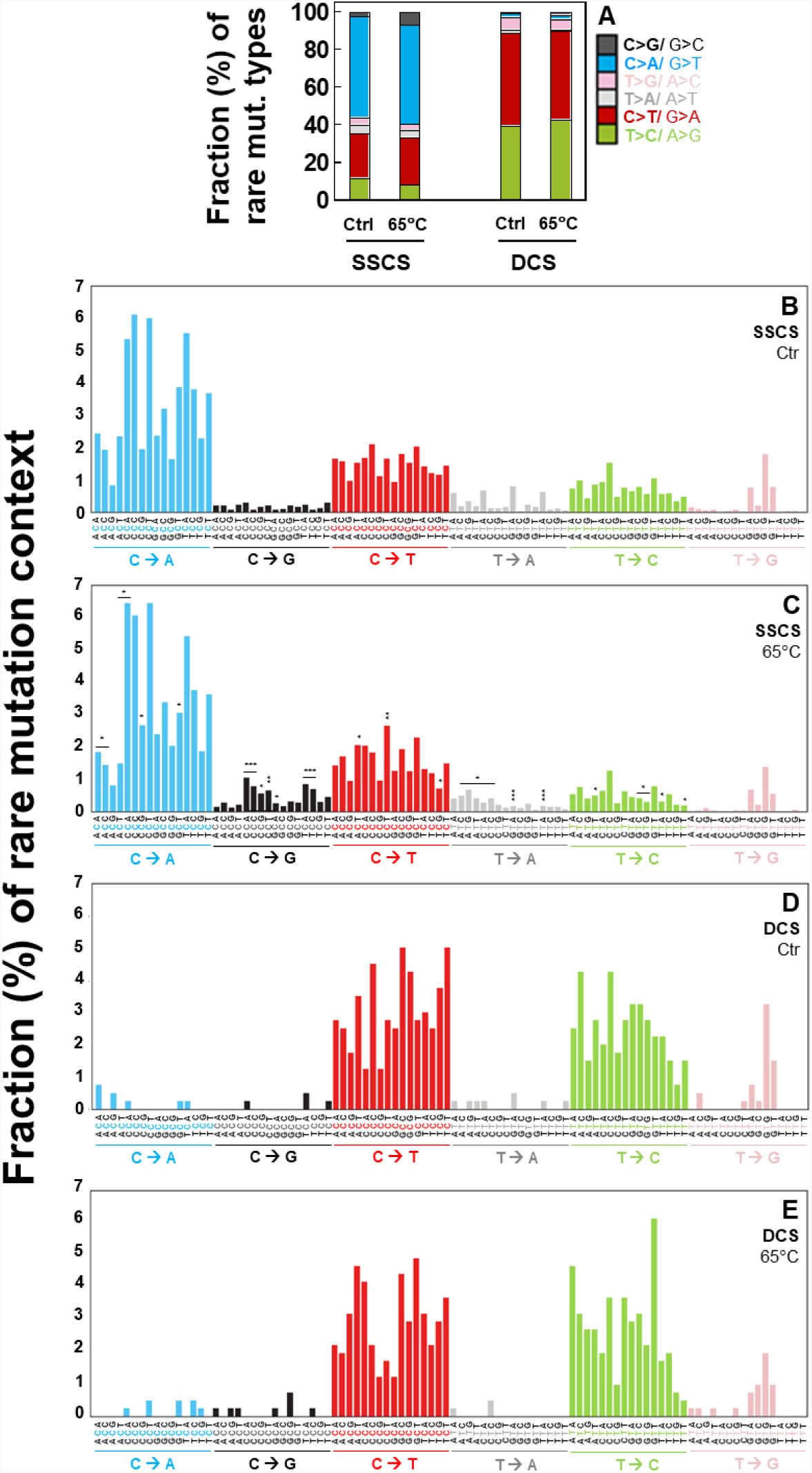
Fractions (%) of the heat-induced (65°C) artifactual mutation types and context spectra in human breast normal cells. **(A)** Relative percentages of rare mutation types for heated versus control DNA as determined by SSCS and DCS analyses. **(B-E)** Rare mutation context spectra for heated versus control DNA as determined by SSCS and DCS analyses. The significant differences in percentage of each mutation context between the control untreated DNA and the heated DNA under SSCS analyses **(C)** are indicated (* *p* < 0.05, ** *p* < 5 x 10^−4^, *** *p* < 5 x 10^−5^ by the Chi-square test).

### 2.1 Homoplasmic Mutations are Detectable by All Three Methods (NGS, Tag-Based Single Strand Sequencing, and Duplex Sequencing) with the Similar Accuracy and Reproducibility

The overall frequencies (Figure 1A) of homoplasmic mutations and frequencies of each mutation type (Figure 1B-D) are almost identical across all independent experiments of NGS (Q30^r^), SSCS, and DCS analyses.

Fractions (%) of each mutation type (Figure 2A) and each mutation context spectrum (Figure 2B-D) were examined for homoplasmic mutations. A mutation context spectra analysis identifies bases immediately 5’ and 3’ to a mutated base (i.e. the mutation appears at the second position of each trinucleotide) and enables classifying observed substitutions into 96 categories (4 bases x 6 substitutions x 4 bases) [13,14].

In our study, fractions (%) of each homoplasmic mutation type (Figure 2A) and their mutation context spectra (Figure 2B-D) are almost same across all the three sequencing methods and independent experiments (Figure 2). All of the homoplasmic mutation types are T>C/A>G or C>T/G>A transitions. T>C/A>G transitions constitute 70% of these mutations and C>T/G>A transitions make up the remaining 30% (Figure 2A). We detected 35 identical homoplasmic unique mutations in all independent experiments regardless of sequencing methods used (Table S3). Taken together, all three sequencing methods are accurate enough to study highly prevalent mutations such as germline mutations of the nuclear genome and homoplasmic mutations of the mitochondrial genome.

### 2.2. Rarely Occurring Mutations are Neither Accurately Detectable by Conventional NGS Methods nor Tag-Based Single Strand DNA Sequencing, but are Accurately Detectable By Duplex Sequencing

Rare mutation frequencies of human breast immortalized cells were determined using NGS, SSCS, and DCS methods. The average rare mutation frequencies of the independent experiments are significantly lower in SSCS (1.30×10^−4^) and DCS (1.04×10^−5^) by 5-fold and 67-fold respectively than that of NGS (Q30^r^) (7.00×10^−4^) (Figure 3A, Table S1). This indicates that Duplex Sequencing removes false-positive artifactual mutations and significantly reduces the rare mutation frequencies.

The frequencies of rare mutations are highly variable in independent experiments analyzed with conventional NGS (Q30^r^) (Figure 3A), whereas frequencies of rare mutations show reproducible results in independent experiments of DCS of Duplex Sequencing (Figure 3A,C). It is noted that NGS (Q30^r^) datasets were processed under more conservative conditions (See section: *Materials and Methods 4.3.2*) than those of SSCS and DCS; however, these bioinformatical modifications only lowered rare mutation frequency by, on average, 35% (Figure S3,S4 and Table S1). This indicates that the bioinformatical modification alone is not possible to overcome the high error rate of NGS.

Frequencies of each type of rare mutations reveal other significant differences between NGS (Q30^r^), SSCS, and DCS analyses. In NGS (Q30^r^) results, C>T/G>A transitions and C>A/G>T transversions are identified at high frequencies (Figure 3D-G). In SSCS results, C>A/G>T transversions are the most predominant mutation type followed by C>T/G>A transitions (Figure 3H,I). In contrast, DCS results indicate that C>T/G>A transitions and C>A/G>T transversions are no longer predominant and no particular type is more prominent than others (Figure 3J,K). Our data suggest that C>A/G>T transversions appear to be the most prevalent type of artifactual mutations that are scored by both NGS (Q30^r^) (Figure 3D-G) and SSCS (Figure 3H,I) methods.

Proportions (%) of each type of rare mutations were analyzed. The prevalent rare mutation types differ under the three sequencing methods. C>A/G>T transversions are the most dominant type of rare mutation with NGS (Q30^r^) data (Figure 4A); however the exact fraction (%) of C>A/G>T transversions vary widely across the four independent NGS experiments. In contrast, comparable fractions (%) of each mutation type are observed in both DCS independent experiments (Figure 4A).

Significant variations are observed for rare mutation context spectra of NGS (Q30^r^) data across four independent experiments (Figure 4B-E) and it is not possible to identify a dominant mutation context among them. In contrast, rare mutation context spectra are similar in all independent DCS experiments. For example, <u>C</u>>T transitions in contexts A<u>C</u>A and A<u>C</u>T occur at persistently high proportions in DCS data (Figure 4H,I).

### 2.3. Duplex Sequencing Identifies and Corrects the Heat-Induced Artifactual Mutations Introduced During DNA Sample Preparation

We investigated which specific types of artifactual mutations are introduced during DNA sample preparation such as heat treatments and to what extent these artifacts can be corrected by Duplex Sequencing. DNA was isolated from human breast primary normal cells (II) and an aliquot of DNA was incubated at 65°C for 9 hours. Unheated DNA served as the control. Libraries of heated and control DNA were prepared for Duplex Sequencing. To identify heat-induced specific mutation types, we performed both SSCS and DCS analyses. The average SSCS and DCS depths of the whole mtDNA genome were similar for control DNA and heated DNA: SSCS (control: 12,257 and heated: 11,622) and DCS (control 2,248 and heated: 2,510) (Table S2). To closely examine the DNA damage-induced artifactual mutations, which are not detectable or distinguishable by conventional sequencing methods, we investigated the rare mutations that include only variants occurring at a frequency of 1% or less using Duplex Sequencing.

The rare mutation frequencies of SSCS are significantly higher than those of DCS (Figure 5A). This higher SSCS mutation frequency could be due to heat-induced DNA damage and/or errors during PCR-amplification. Such artifactual mutations are present on only one of the two DNA strands and thus they are not scored in DCS of Duplex Sequencing. While the incubation of DNA at 65°C significantly increased rare mutation frequency in SSCS analysis (Figure 5A: the first and second bars: *p*-value < 2.2×10^−16^), both heated and control DNA displayed identical frequencies of rare mutations in DCS analysis (Figure 5A: the third and fourth bars). Our results clearly indicate that DCS analysis by Duplex Sequencing is not affected by heat-induced DNA damage introduced during DNA sample preparation and correctly represents true mutations.

### 2.4. Duplex Sequencing Identifies the Specific Mutation Spectra of Heat-Induced Artifacts

We further examined which specific mutation type(s) contributed to the elevated SSCS rare mutation frequency in heated DNA. In SSCS, but not DCS, the heated DNA (Figure 5B) shows a significant increase in rare mutation frequencies of C>A/G>T, C>T/G>A, C>G/G>C *versus* control DNA (Figure 5B). In contrast, the 65°C incubation (heating DNA) did not affect the mutation spectra of DCS results (Figure 5C). For example, the SSCS mutation frequency of C>A in heated DNA is 3.26×10^−5^. This heat-induced artifactual mutation type is significantly reduced by 170-fold to 1.88 × 10^−7^ in DCS analysis.

Fractions (%) of each type (Figure 6A) and each context spectrum of rare mutations (Figure 6B-E) were examined for heated *vs.* control DNA. The heat-induced DNA damage results in increases in C>G/G>C in SSCS analysis (Figure 6A-C). Out of the 96 possible mutation sequence contexts, 28 are significantly changed after the 65°C incubation in SSCS analysis (Figure 6B-C, Table S4). Particularly, C<u>C</u>C, T<u>C</u>C and C<u>C</u>A contexts of <u>C</u>>G mutations showed the most significant increase in the heated DNA compared to the control (unheated) DNA in SSCS analysis (Figure 6C, Table S4). In contrast, these DNA damage-dependent substitutions are not observed in DCS analysis, irrespective of neighboring nucleotides (Figure 6A,D,E).

### 2.5. Independent Experiments of Duplex Sequencing Reproducibly Identify the Heat-Induced Artifactual Mutaiton Profiles

Two independent experiments (I and II) for the incubation of DNA at 65°C for 9 hours and the DNA library preparation was conducted with DNA isolated from human epithelial cells (I and II) derived from breast tissue of the same woman. The average SSCS and DCS depths were similar between the cells I and cells II: SSCS (I: 11,622 and II: 11,045) and DCS (I: 2,510 and II: 2,460) (Table S5).

Rare mutation frequencies of heated DNA of the cells I (Figure S5) were calculated for both SSCS and DCS, and the analysis reveals the same pattern observed with heated DNA of the cells II used for the main result Figure 5 experiment (Figure S5). Both the overall rare mutation frequencies and the rare mutation frequency of each mutation type are observed at similar levels between the heated DNA of the cells I and II (Figure S5). Furthermore, fractions (%) of each mutation type (Figure S6A) and mutation context spectra (Figure S6B-E) are almost identical, strengthening the finding that Duplex Sequencing is capable of identifying and correcting the heat-induced artifactual mutations and the results are reproducible in independent experiments.

### 2.6. All Identified Heat-Induced Artifactual Mutations are Stochastically Occurring Mutations Throughout the Whole Mitochondrial Genome

A total of 3,383 heat-induced artifactual unique mutations were identified, all these are in the mutation occurrence (%) range of 0-1% (Figure 7A). This clearly indicates that all of the heat-induced artifactual mutations introduced during DNA preparation are rarely occurring mutations, thus not accurately and reliably detectable by conventional DNA sequencing methods.

**Figure 7.**
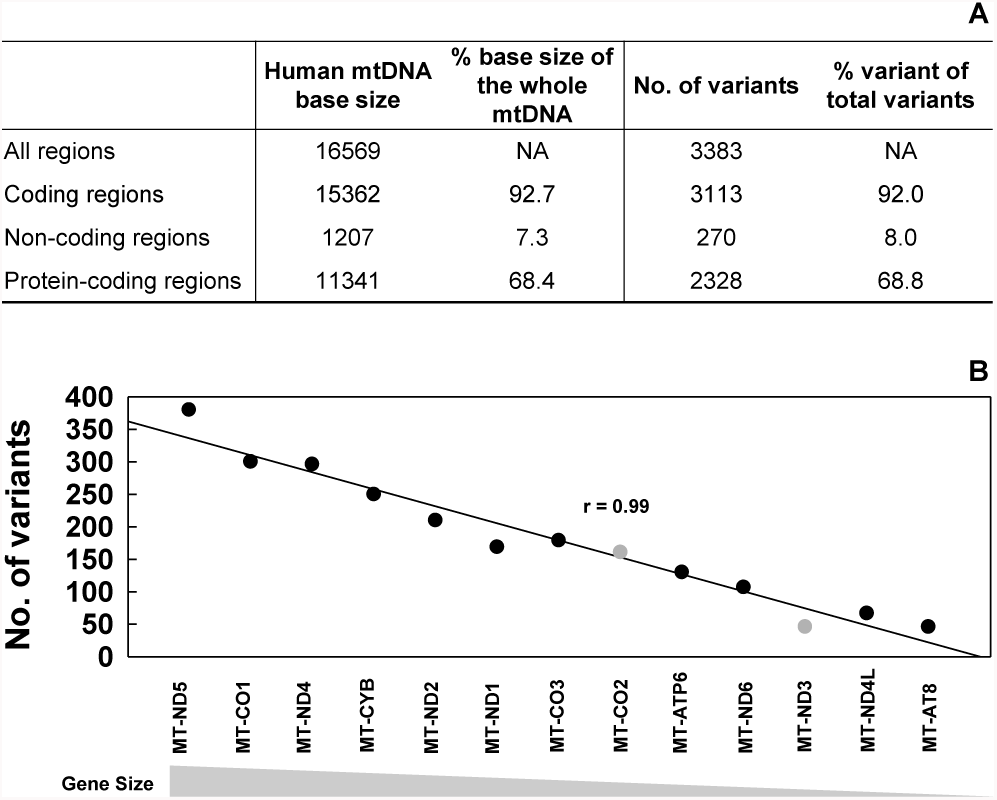
Numbers of the heat-induced (65°C) artifactual mutations in the whole mtDNA of human breast normal cells identified by SSCS and DCS analyses. Variants were counted only once at each position of the genome. (**A**) Numbers of variants identified in coding, protein-coding, and non-coding regions of mtDNA. The percentage of variants from each region out of the total number of identified variants is calculated (% variants). The percentage of base size that each region occupies out of the total size of mtDNA is calculated (% base size). (**B**) The numbers of the heat-induced artifactual rare mutations identified in each of 13 protein-coding genes of mtDNA are plotted in the order of largest to smallest gene size. The Pearson’s correlation coefficient for each gene size of the 13 genes *versus* numbers of variants of each gene was 0.99 (*p* = 2.36 x 10^−10^).

Of the all identified artifactual mutations, about 92% of them are found on coding regions of mtDNA and 69% of them are found within the 13 protein-coding regions of mtDNA (Figure 7A, Table S6). The percentages of artifactual mutations found on coding regions and 13 protein-coding regions of mtDNA closely match the percentages these two regions occupy in the whole mtDNA, which are about 92% and 68% respectively [15]. In addition, the number of artifactual mutations identified shows a strong positive correlation with the sizes of 13 protein-coding genes (Pearson’s correlation coefficient R=0.99 and *p*=2.36×10^−10^), which indicates that more artifactual mutations are found on larger genes (Figure 7B). We examined the 13 protein-coding genes of mtDNA to see if any particular gene is relatively more or less prone to heat-induced mutations. For each gene, we calculated the percent of variants by dividing the numbers of variants by each gene size (bases). Among the 13 genes, MT-CO2 is slightly more mutated at 23.39% and MT-ND3 is mutated the least at 13.01% (Table S7). The majority (11 out of 13) of these genes are mutated at similar mutation occurrences (average 20%), which is consistent with the finding that artifactual mutations occur stochastically.

## 3. Discussion

In this study, we sequenced the entire mitochondrial DNA of human breast cells via three different sequencing protocols: NGS, tag-based single strand sequencing (eg. SSCS), and Duplex Sequencing. We systematically compared high-frequency and low-frequency mutations obtained from the three methods. We demonstrate advantages of Duplex Sequencing over other sequencing methods for studying rarely occurring mutations. In addition, we identified the heat-induced artifactual mutations. Although Duplex Sequencing has been used in a previous study to show an increased level of mutation frequency of small selected regions of nuclear genome in DNA incubated at 65°C [11], the same temperature used in this study, exact identities of the heat-induced artifactual mutations have not been presented. Moreover, while types of artifactual mutations have been previously examined, influences of neighboring nucleotide base context on artifactual mutations have not been investigated. To our knowledge, this is the first study to present the exact identities of the heat-induced artifactual mutations and the specific nucleotide context spectra for these artifacts.

Our data show that rare mutation frequencies are significantly lower in DCS analysis in comparison to NGS and SSCS analyses, suggesting that large number of rare mutations detected by NGS and SSCS are mostly artifacts (Figure 3A). Particularly, C>A/G>T transversions, which have been previously reported to be a predominant result of DNA oxidation [11,16], showed the greatest decrease in frequencies with DCS analyses of Duplex Sequencing (Figure 3J,K). These findings validate that DCS can identify and correct artifacts and be applied for accurately detecting rarely occurring mutations. Furthermore, the comparison of rare mutation frequencies between multiple independent experiments (Figure 3) demonstrates the ability of DCS in producing reproducible results. However, the rare mutation data by NGS and SSCS shows high variability across independent experiments, which indicates the lack of the capability of NGS and SSCS to produce reliable and reproducible results. The comparison of rare mutation context spectra analyses (Figure 4) further distinguishes DCS from NGS and SSCS by showcasing its advantage to accurately and consistently detect rarely occurring mutations.

Artifactual mutations can be generated as a result of copying damaged DNA bases. Such mutations are present on only one of the two DNA strands and are scored by NGS and SSCS but not by DCS. In the present study, we have identified heat-induced artifactual mutations by performing both SSCS and DCS analyses. Our results indicate that C>A/G>T is the most predominantly enhanced mutation artifact followed by C>T/G>A and C>G/G>C. Previous studies reported that heating DNA can damage DNA bases by forming oxygen free radicals, (specifically 8-hydroxy-2’-deoxyguanine (8-Oxo-dG)), which deaminate cytosine to uracil, and increasing mitochondrial superoxide anion, which also leads to oxidation of DNA [17–19]. The 8-Oxo-dG is generated by DNA oxidation under physiopathological conditions or environmental stress. It is also a by-product of normal cellular metabolism [20]. The formation of 8-oxoguanine, particularly 8-oxo-dG has been reported to cause a high level of C>A/G>T mutations [11,16,20–23], whereas deamination of cytosine to uracil is known to produce high levels of C>T/G>A mutations [11,19]. In one study, NGS analysis was done on tumors and matching normal tissues of melanoma and an enzyme-linked immunosorbent assay (ELISA) for 8-Oxo-dG found that the C<u>C</u>G>C<u>A</u>G context have a high potential for being a target of DNA oxidation [22]. This context is observed at high mutation frequencies (2-fold increase) in our current study. Thus, it is likely that the high prevalence of C>A/G>T transversions in our data is caused by 8-Oxo-dG, suggesting that the biggest contributor of heat-induced artifactual mutations is oxidative damage to mtDNA.

To our knowledge, this study is the first to examine mutation context spectra of heat-induced artifactual mutations in the whole mitochondrial DNA of human breast epithelial cells. Our SSCS analysis results showed that out of 96 possible mutation sequence contexts, the fraction of 28 rare mutation sequence contexts were significantly changed after the 65°C incubation (Figure 6B-C, Table S4). Among the affected 28 mutation context spectra, C<u>C</u>C, T<u>C</u>C and C<u>C</u>A contexts of <u>C</u>>G mutations showed the most significant increase. These mutation contexts could be more prone to DNA damage and may have a high potential for being targeted by molecular reactions which result from the heat-damaged DNA.

In summary, we present a genome-wide comprehensive and comparative analysis of mitochondrial DNA mutations for NGS and tag-based methods (single-strand sequencing and Duplex Sequencing) and demonstrate the identification and removal of heat-induced artifactual mutations using Duplex Sequencing. Our results indicate that all of the heat-induced artifactual mutations are stochastically occurring rare mutations. Thus, these artifactual mutations are not accurately detectable by conventional sequencing methods. Even the application of more conservative bioinformatical modification on NGS datasets is not enough to overcome the inherently high error rate of conventional NGS methods. Our data establishes that Duplex Sequencing: 1) accurately and reproducibly detect rare (low-frequency) mutations; 2) is not affected by damage introduced by heating during DNA preparation; 3) identifies and removes the DNA damage-induced artifactual mutations.

## 4. Materials and Methods

### 4.1. Cell Culture

Human breast epithelial cells were provided by Drs. Chia-Cheng Chang and James E. Trosko at Michigan State University in East Lansing, MI, USA. Human breast normal cells used in this study were isolated from breast tissues of a healthy (cancer-free) woman 21 years of age obtained during reduction mammoplasty at Sparrow Hospital in Lansing, MI, USA. Human breast immortalized cells (M13SV1) were derived from the parental normal stem cells by transforming with SV40 T antigen [24–29]. Written consents were received from patients. The use of human breast cells was approved by Michigan State University Institutional Review Board and a Material Transfer Agreement was approved by both Michigan State University and University of Washington. The cells were cultured as previously described [30] and were authenticated by short tandem repeat (STR) DNA profiling (Genetica DNA Laboratories, Labcorp brand, Burlington, NC, USA).

### 4.2. DNA Extraction, Adapter Synthesis, Library Preparation

DNA extraction and purification, adapter synthesis, and DNA library preparation for Duplex Sequencing [31] and for whole exome sequencing (WES) [32,33] were carried out as described previously. For Duplex Sequencing, DNA was extracted using a commercially available DNA extraction kit (Invitrogen) and the sheared DNA was subjected to end-repair with 3’-dT-tailing for adapters with A-overhang and with 3’-dA-tailing for adapter with T-overhang. The whole mitochondrial genome was captured using Agilent SureSelect^XT^ target enrichment (Agilent Technologies). For WES, the DNA library was hybridized to biotinylated capture probes from the SureSelect Human All Exon kit (Agilent Technologies). This kit covers 38 Mb of human genome, corresponding to 23,739 genes in the National Center for Biotechnology Information Consensus Coding Sequence database.

### 4.3. Data Processing

#### 4.3.1 NGS Datasets and NGS, SSCS and DCS Data Processing

Two of four independent NGS datasets (Experiments #1 and #2 in Figure 1 through Figure 4) were obtained by extracting the whole mitochondrial genome data from our WES results. WES is a commonly used next-generation sequencing method that is able to read the entire exome of DNA as well as the mitochondrial genome [34]. The fastq data files for the two WES datasets were processed as previously described [33] with some modifications. Our in-house script was modified to align the reads with the human mitochondria reference file and to include the mitochondrial genome only.

Two additional independent NGS datasets were obtained by modifying our in-house Duplex Sequencing script to simulate NGS processing on DNA libraries prepared for Duplex Sequencing (Experiments #3 and #4 in Figure 1 through Figure 4). The modified script proceeds only through alignment with a human mtDNA reference but does neither single strand consensus sequence (SSCS) nor duplex consensus sequence (DCS) data alignment steps. This negates the individuality of complementary DNA strands when processing sequence data.

SSCS and DCS datasets were processed as described previously [31]. All datasets were aligned to the Revised Cambridge Reference sequence (rCRS) reference genome, sequence number NC_012920, using BWA and genome analysis toolkit (GATK) software as described previously [31].

#### 4.3.2. Base Quality and PCR Duplicates

An illumina^®^ base quality score of 30 (Q30) is considered the benchmark for a correct base call in NGS. This score refers to a 1 in 1000 chance of an incorrect base call (error probability of 0.1% *vs.* error probability of 5% with the default score of 13) [35] Our NGS (Q30^r^) datasets, therefore, were processed with the base quality filtering adjusted to 30 from the default value of 13 by adding “-Q30” to the pileup command (i.e. samtools mpileup –B –d 500000 –Q30 –f [reference] [input] [output]). However, introduction of artifacts in early stages of PCR amplification are not detectable as errors and are embedded in multiple PCR duplicates [6]. SAMtools software is capable of removing these PCR duplicates [36], and so they were removed for NGS (Q30^r^) data by taking the combined sequence-1 and sequence-2 files in bam format and using the command “samtools rmdup –s”. Both modifications bioinformatically accommodate high background error rates of conventional NGS. For DCS and SSCS data analyses, the default base quality score of 13 (error probability of 5%) was used and PCR duplicate removal was not applied since SSCS and DCS analyses have significantly lower error frequencies (1×10^−5^ and <5×10^−8^ or 1×10^−8^, respectively) than that of conventional NGS (10^−2^ to 10^−3^) [4,9]. Another reason the PCR duplicate removal step is not needed for SSCS and DCS analyses is that the molecular tags mark the duplicates for SSCS and DCS analyses.

The results of NGS analysis obtained under the same bioinformatical conditions as SSCS and DCS (Q13 and no PCR duplicate removal) were presented in Table S1. The effects of bioinformatical modifications (Q13 and no PCR duplicate removal *versus* Q30 and PCR duplicate removal) on NGS mutation results are presented in Supplementary Figures 1-4.

#### 4.3.3. Comparison of Mutation Positions

To identify the heat-induced artifactual mutations, mutation positions were compared after removing the common mutations (present in both control and heated DNA) identified by SSCS and DCS analyses. Mutations found only in the heated DNA from SSCS analysis were considered as heat-induced artifactual mutations. Only genome positions that had minimum sequence read (depth) of 20 in both samples were considered.

#### 4.3.4. Counting mutations

For calculating mutation frequencies, total number of variant reads observed were divided by total number of sequenced reads. For all other analyses, including fractions (%) of each mutation type and mutation context spectra, mutants were scored only once at each position of the genome (i.e. mutants were counted as 1 for each position regardless of number of variant reads observed in that position).

### 4.4. Statistical Analysis

Differences in mutation frequencies and mutation context fractions between control and heated DNA were analyzed by performing a Chi-square test using R (program version 3.4.4). Association of the number of identified unique variants in each of the thirteen protein-coding genes of mtDNA and the sizes of the corresponding thirteen genes was analyzed by Pearson correlation using Sigma Plot (version 12.0, Systat Software, Inc., San Jose, CA, USA). Differences between the groups were considered significant at *p*<0.05.

## Supporting information

## Author contributions

EHA conceived and designed the experiment; EHA performed the experiments; SHL and EHA processed sequencing data and analyzed the data; EHA and SHL wrote the paper.

## Acknowledgements

The research was supported by grants from the National Institute of Environmental Health Sciences (NIEHS) P30 ES007033 sponsored-University of Washington (UW) Center for Exposures, Diseases, Genomics and Environment (EDGE) pilot grant (to EH Ahn), UW Office of Research Royalty Research Fund (to EH Ahn), and NCI P30 CA015704-39 Fred Hutchinson Cancer Research Center-UW Cancer Consortium Support Grant (to EH Ahn), National Cancer Institute (NCI) R21 CA220111 (to EH Ahn) and National Cancer Institute of the National Institutes of Health under award number P01 AG001751 and R33 CA181771 (to LA Loeb). The content is solely the responsibility of the authors, and does not necessarily represent the official views of the National Institutes of Health. We thank Dr. Lawrence A. Loeb for critical reading of this manuscript, Dr. Tom Walsh for NGS technical assistance, Drs. Chia-Cheng Chang and James E. Trosko for providing human breast cells, Howard Nebeck for assistance in mutation context analysis and proofreading the manuscript, Clint Valentine, Dr. Michael W. Schmitt, Dr. Woo Young Kim for bioinformatics consultation, and Sujin Kwon for graphical assistance.

## Abbreviations

DCS: Duplex Consensus Sequence
DS: Duplex Sequencing
MT: Mitochondria
NGS: Next-generation Sequencing
SSCS: Single Strand Consensus Sequence

