## Supplementary material for "Detection of low-frequency mutations and removal of heat-induced artifactual mutations using Duplex Sequencing"

**Table S1.** Sequencing data yield of human breast immortalized cells. The average depth (number of nucleotides sequenced at each genome position) was calculated as the total number of nucleotides sequenced divided by the mitochondrial (mt) DNA size of 16569 bases.

**A.** NGS and NGS (Q30$^r$) data yield (Data used for Figures 1-4 and S1-S4)

| Exp | Avg NGS depth | Total sequenced NGS Nts | **NGS: No. of rare muts** | **NGS: Rare mut freq** | Avg NGS (Q30$^r$) depth | Total sequenced NGS (Q30$^r$) Nts | **NGS (Q30$^r$): No. of rare muts** | **NGS (Q30$^r$): Rare mut freq** |
|---|---|---|---|---|---|---|---|---|
| #1 | 517 | 8558610 | **9803** | **1.15 x10$^{-3}$** | 109 | 1810307 | **1131** | **6.25 x10$^{-4}$** |
| #2 | 2130 | 35293720 | **45526** | **1.29 x10$^{-3}$** | 138 | 2279999 | **556** | **2.44 x10$^{-4}$** |
| #3 | 210473 | 3487319586 | **2476099** | **7.10 x10$^{-4}$** | 140 | 2311906 | **2395** | **1.04 x10$^{-3}$** |
| #4 | 458441 | 7595916093 | **8773866** | **1.16 x10$^{-3}$** | 140 | 2322756 | **2084** | **8.97 x10$^{-4}$** |

**B.** SSCS and DCS data yield (Data used for Figures 1-4 and S1-S4)

| Exp | Avg SSCS depth | Total sequenced SSCS Nts | **SSCS: No. of rare muts** | **SSCS: Rare mut freq** | Avg DCS depth | Total sequenced DCS Nts | **DCS: No. of rare muts** | **DCS: Rare mut freq** |
|---|---|---|---|---|---|---|---|---|
| #3 | 2086 | 34562417 | **3838** | **1.11 x10$^{-4}$** | 444 | 7363685 | **77** | **1.05 x10$^{-5}$** |
| #4 | 40421 | 669733398 | **99361** | **1.48 x10$^{-4}$** | 6803 | 112714571 | **1164** | **1.03 x10$^{-5}$** |

Abbreviations used are: Avg, average; Nts, nucleotides; mut, mutation; freq, frequency

**Table S2.** Sequencing data yield for heated *vs.* control DNA of human breast normal cells (II).

| Cells | Avg SSCS depth | Total sequenced SSCS Nts | **SSCS: No. of rare muts** | **SSCS: Rare mut freq** | Avg DCS depth | Total sequenced DCS Nts | **DCS: No. of rare muts** | **DCS: Rare mut freq** |
|---|---|---|---|---|---|---|---|---|
| Control (Data for Figs 5, 6) | 12257 | 203078000 | **9351** | **4.60 $\times 10^{-5}$** | 2248 | 37240836 | **577** | **1.55 $\times 10^{-5}$** |
| Heated (Data for Figs 5, 6, S5, S6) | 11622 | 192572847 | **15429** | **8.01 $\times 10^{-5}$** | 2510 | 41582269 | **635** | **1.53 $\times 10^{-5}$** |

Abbreviations used are: Avg, average; Nts, nucleotides; SSCS, single strand consensus sequences; DCS, duplex consensus sequences; mt, mitochondria; mut, mutation; freq, frequency

**Table S3.** Identical homoplasmic unique mutations detected using next-generation sequencing (NGS Q30[r]), SSCS, and DCS analyses in all independent experiments for mtDNA of human breast immortalized cells. The mutations are listed in the order of largest to smallest gene size. Green represents T>C/A>G mutations and red represents C>T/G>A mutations.

| **Gene** | **DNA mutation** | **Amino acid change** | **Mutation type** |
|---|---|---|---|
| MT-ND5 | G12372A | *Syn* | [red] |
| | A13117G | I261V | [green] |
| MT-RNR2 | A1811G | *Syn* | [green] |
| | A2706G | N346D | [green] |
| MT-CO1 | G6179A | *Syn* | [red] |
| | C6587T | *Syn* | [red] |
| | C7028T | *Syn* | [red] |
| MT-ND4 | T11299C | *Syn* | [green] |
| | A11362G | *Syn* | [green] |
| | A11467G | *Syn* | [green] |
| | G11719A | *Syn* | [red] |
| MT-CYB | C14766T | T7I | [red] |
| | T14798C | F18L | [green] |
| | A15326G | T194A | [green] |
| MT-ND1 | A3480G | *Syn* | [green] |
| | A4769G | *Syn* | [green] |
| MT-RNR1 | A750G | N35D | [green] |
| | T1189C | I181T | [green] |
| | A1438G | Ter264W | [green] |
| MT-CO3 | T9698C | *Syn* | [green] |
| MT-ATP6 | T8632C | Y36H | [green] |
| | G9055A | A177T | [red] |
| MT-ND6 | C14167T | Syn | [red] |
| MT-ND3 | A10398G | T114A | [green] |
| MT-ND4L | A10550G | *Syn* | [green] |
| MT-TL2 | A12308G | K15E | [green] |
| MT-TD | A7559G | *Syn* | [green] |
| Non-coding | A73G | | [green] |
| | A263G | | [green] |
| | C497T | | [green] |
| | T16093C | | [red] |
| | T16224C | | [green] |
| | T16311C | | [green] |
| | G16390A | | [green] |
| | T16519C | | [red] |

**Table S4.** Significant differences in mutation context counts between heated *vs.* control DNA of human breast normal cells determined using SSCS analysis. Significant $p$ values are denoted with * (<0.05), ** (<$5x10^{-4}$), and *** (<$5x10^{-5}$) by the Chi-square test.

| **C>A** | **C>G** | **C>T** | **T>A** | **T>C** | **T>G** |
|---|---|---|---|---|---|
| ACA * | CCA *** | ACT * | ATC * | ATT * | |
| ACC * | CCC *** | CCT ** | ATG * | GTC * | |
| ACT ** | CCG * | TCG * | ATT * | GTG * | |
| CCA * | CCT ** | | CTA * | TTA * | |
| CCG * | GCA * | | CTC * | TTT * | |
| GCT * | TCA *** | | GTA *** | | |
| | TCC *** | | TTA *** | | |

**Table S5.** Sequencing data yield of human breast normal cells (I and II).

| Cells | Avg SSCS depth | Total sequenced SSCS Nts | **SSCS: No. of rare muts** | **SSCS: Rare mut freq** | Avg DCS depth | Total sequenced DCS Nts | **DCS: No. of rare muts** | **DCS: Rare mut freq** |
|---|---|---|---|---|---|---|---|---|
| II (Data for Figs 5, 6, S5, S6) | 11622 | 192572847 | **15429** | **8.01 $x10^{-5}$** | 2510 | 41582269 | **635** | **1.53 $x10^{-5}$** |
| I (Data for Figs S5, S6) | 11045 | 183007737 | **14881** | **8.13 $x10^{-5}$** | 2460 | 40754407 | **571** | **1.40 $x10^{-5}$** |

Abbreviations used are: Avg, average; Nts, nucleotides; SSCS, single strand consensus sequences; DCS, duplex consensus sequences; mt, mitochondria; mut, mutation; freq, frequency

**Table S7.** Heat-induced artifactual rare mutations identified for the thirteen protein-coding mtDNA genes of human breast normal cells. Variants were counted only once at each position of the genome. The percent variant of each gene was calculated by dividing number of variants observed by the DNA base length of the corresponding gene.

| Protein-coding genes | Gene base length | No. of variants | % variant of each gene |
|---|---|---|---|
| MT-ND5 | 1812 | 379 | 20.92 |
| MT-CO1 | 1542 | 299 | 19.39 |
| MT-ND4 | 1378 | 295 | 21.41 |
| MT-CYB | 1141 | 249 | 21.82 |
| MT-ND2 | 1042 | 209 | 20.06 |
| MT-ND1 | 956 | 168 | 17.57 |
| MT-CO3 | 784 | 178 | 22.70 |
| **MT-CO2** | 684 | 160 | **23.39** |
| MT-ATP6 | 681 | 129 | 18.94 |
| MT-ND6 | 525 | 106 | 20.19 |
| **MT-ND3** | 346 | 45 | **13.01** |
| MT-ND4L | 297 | 66 | 22.22 |
| MT-ATP8 | 207 | 45 | 21.74 |

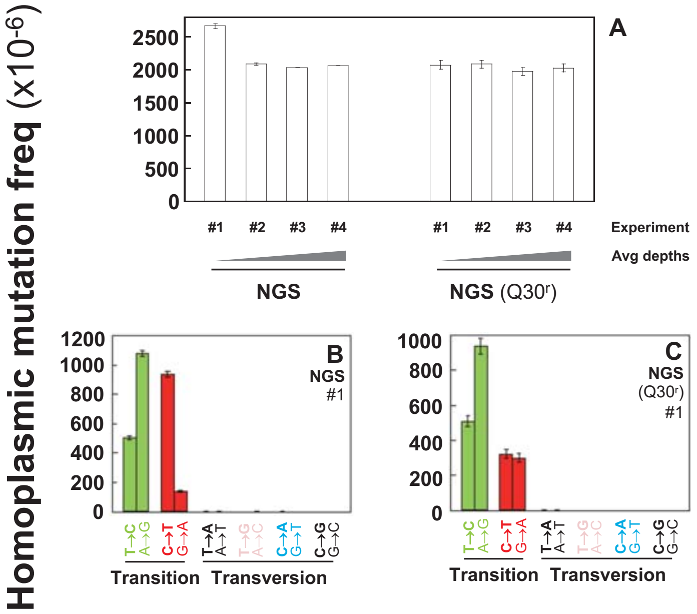


**Figure S1.** Frequencies of homoplasmic point mutations in the whole mtDNA of human breast immortalized cells. (**A**) Overall frequencies determined by performing NGS before and after bioinformatical modifications. The modifications include an increased base quality score of 30 (Q30) from the default score of 13 (Q13) and removal of PCR duplicates. **(B,C)** Homoplasmic mutation frequency of specific mutation types determined using NGS analysis before (B) and after (C) modifications. Error bars represent the Wilson score 95% confidence intervals.

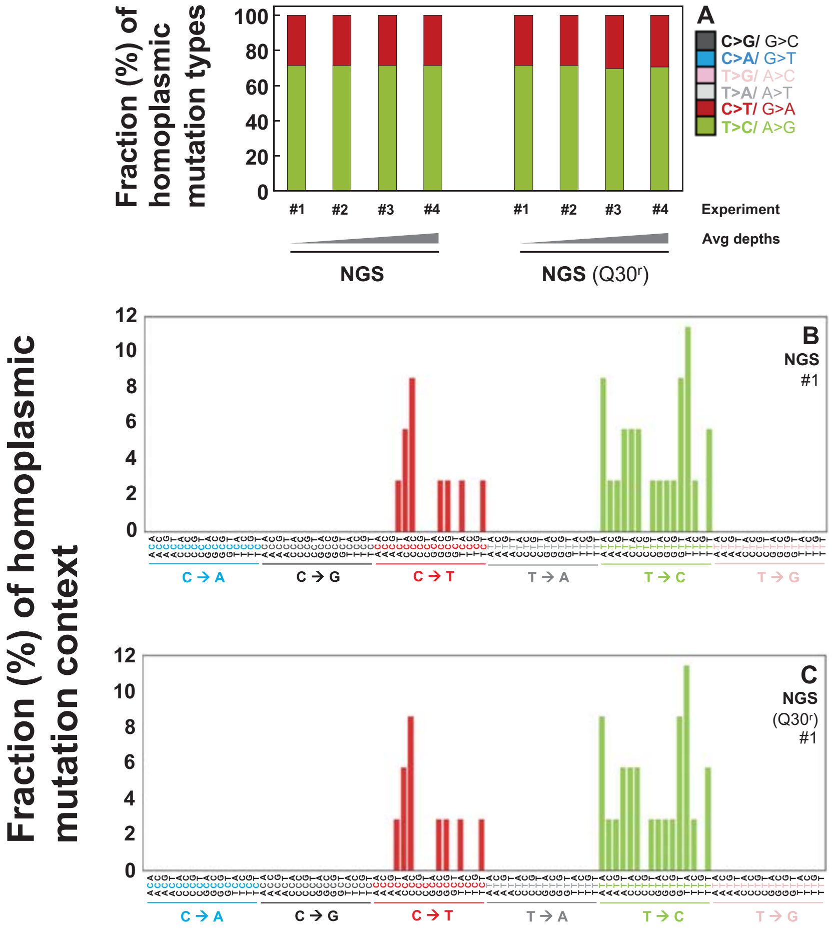


**Figure S2.** Fractions (%) of homoplasmic point mutation types and context spectra in the whole mtDNA of human breast immortalized cells. **(A)** Relative percentages of each mutation type in each experiment as determined by NGS analysis before and after bioinformatical modifications. **(B,C)** Fractions of homoplasmic mutation context spectra determined by performing NGS analysis before (B) and after (C) bioinformatical modifications. The modifications include an increased base quality score of 30 (Q30) from the default score of 13 (Q13) and removal of PCR duplicates.

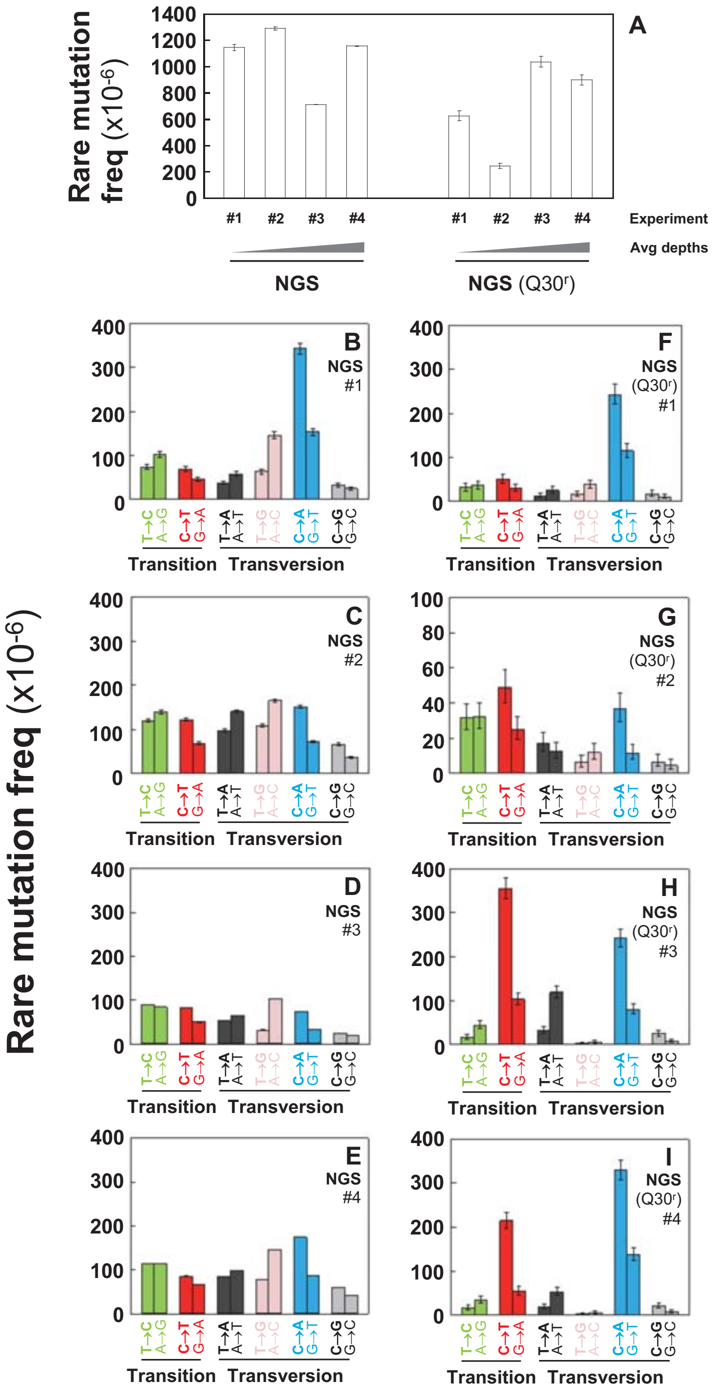


**Figure S3.** Frequencies of rare point mutations in the whole mtDNA of human breast immortalized cells. (**A**) Overall frequencies determined by performing NGS analysis before and after bioinformatical modifications. The modifications include an increased base quality score of 30 (Q30) from the default score of 13 (Q13) and removal of PCR duplicates. **(B-I)** Rare mutation frequency of each mutation type as determined using NGS analysis before (B-E) and after (F-I) modifications. Error bars represent the Wilson score 95% confidence intervals.

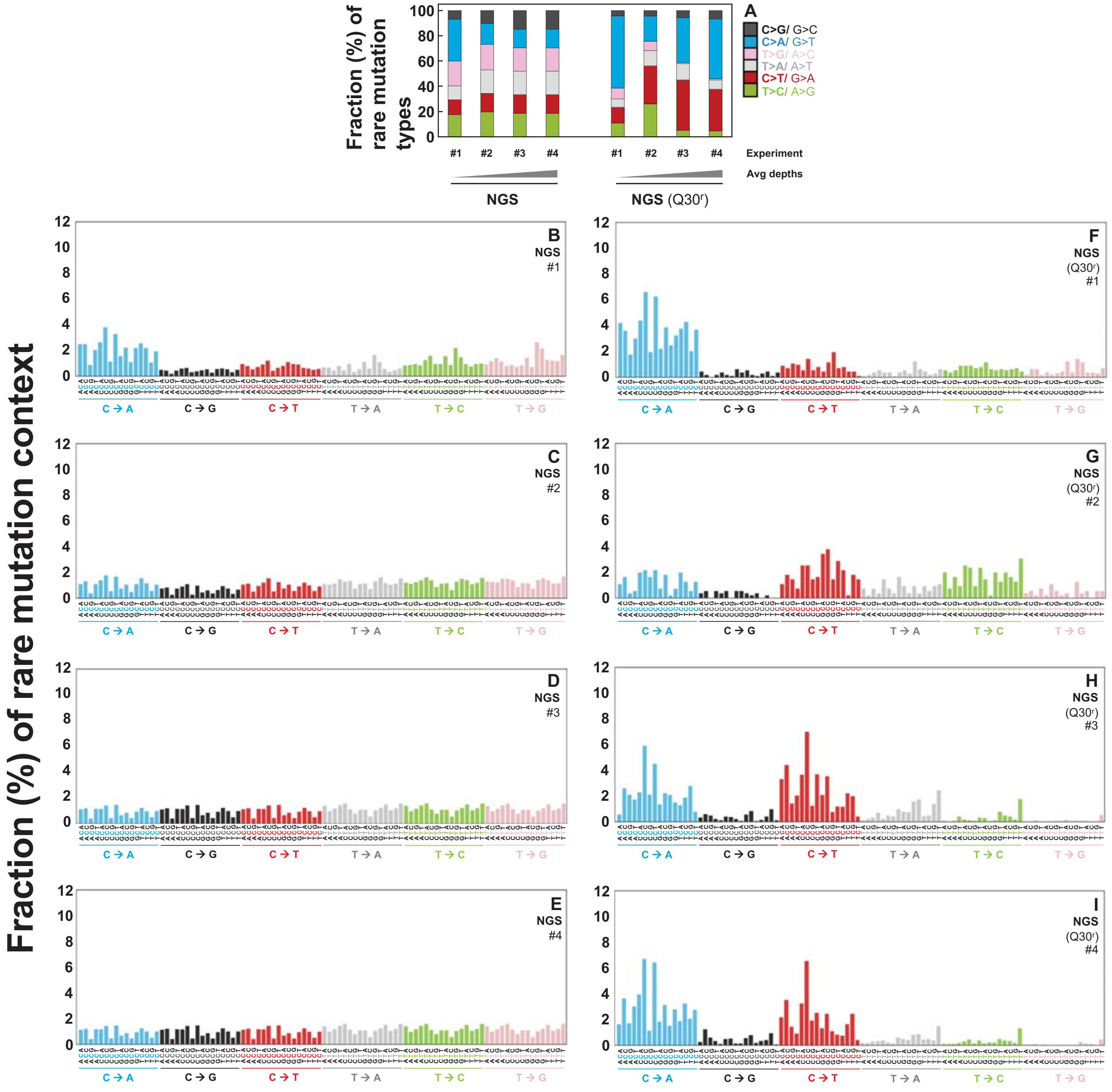


**Figure S4.** Fractions (%) of rare mutation types and context spectra in the whole mtDNA of human breast immortalized cells. (**A**) Relative percentages (%) of rare mutation types in each experiment as determined by performing NGS analysis before and after bioinformatical modifications. **(B-I)** Fractions of rare mutation context spectra determined by performing NGS analysis before (B-E) and after (F-I) bioinformatical modifications. The modifications include an increased base quality score of 30 (Q30) from the default score of 13 (Q13) and removal of PCR duplicates.

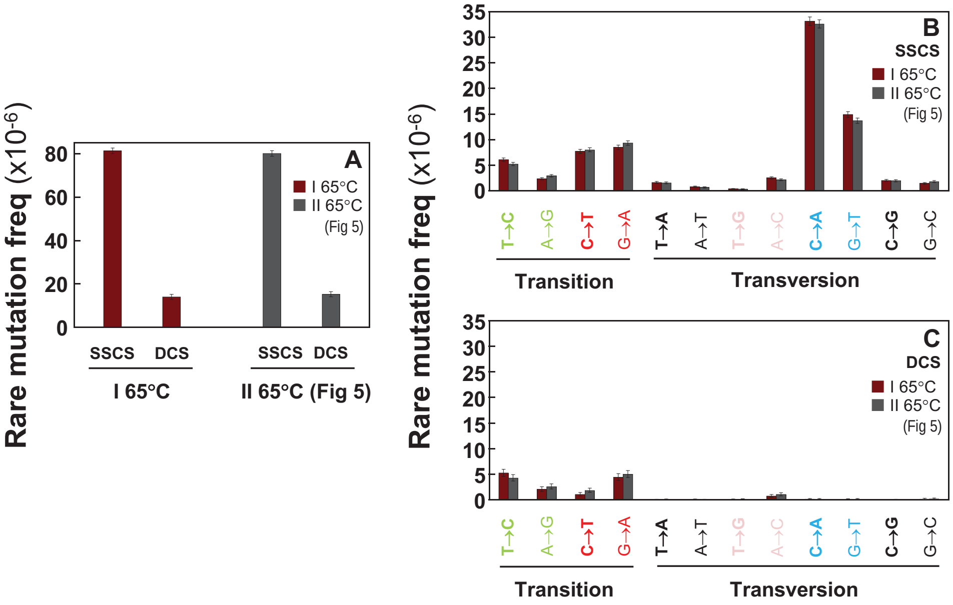


**Figure S5.** Frequencies of the heat-induced (65°C) artifactual rare mutations of human breast normal cells I and II derived from breast tissue of the same woman. Data were generated by conducting two independent DNA library experiments (I and II). The purified DNA were heated at 65°C as described for Fig 5 and Fig 6. Overall rare mutation frequency (**A**) and frequencies of rare mutation types (**B,C**) of the cells were determined by SSCS and DCS analyses. Error bars represent the Wilson score 95% confidence intervals.

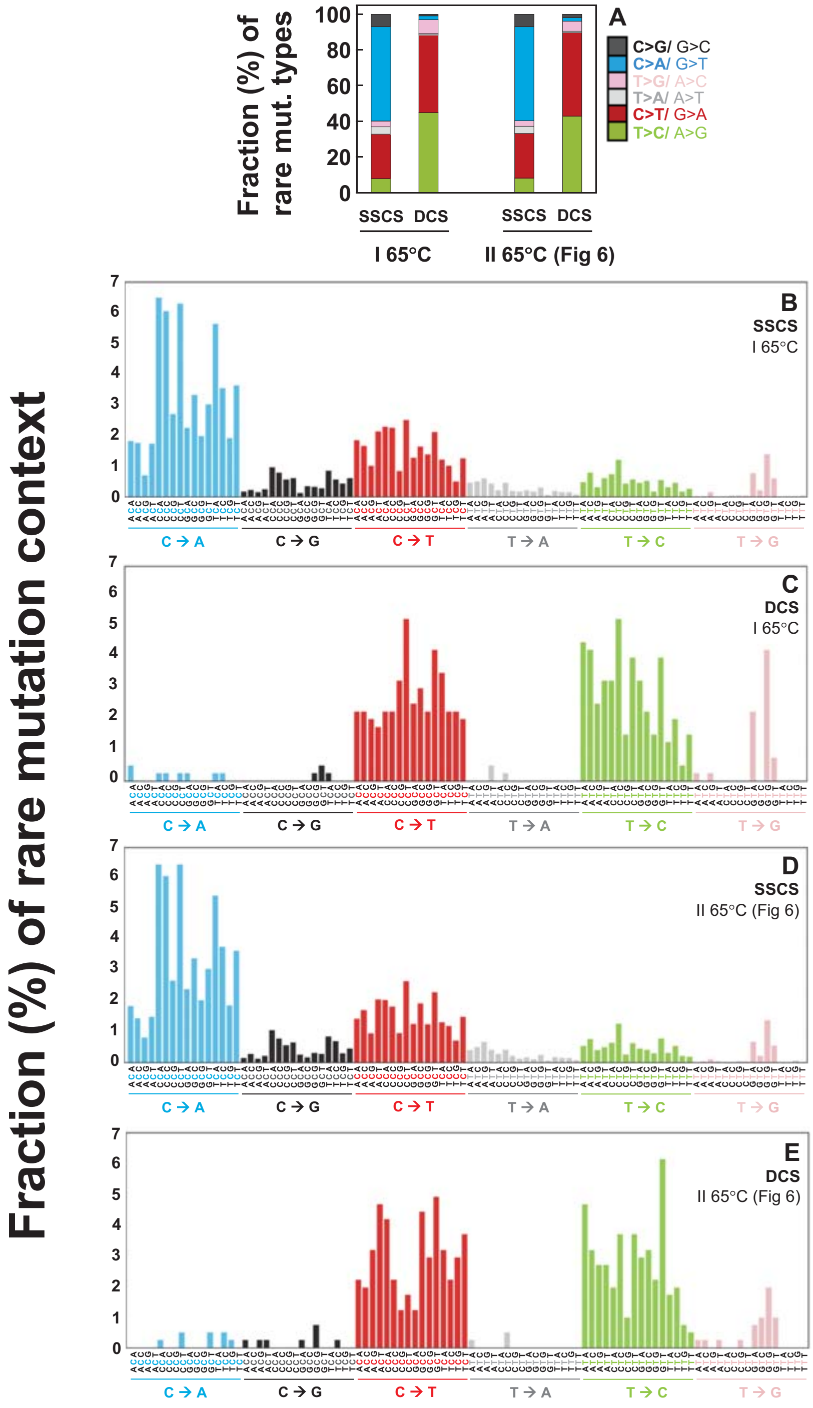


**Figure S6.** Fractions (%) of the heat-induced (65°C) artifactual rare mutation types and context spectra in human breast normal cells I and II derived from breast tissue of the same woman. Data were generated by conducting two independent DNA library experiments (I and II). The purified DNA was heated at 65°C as described for Fig 5 and Fig 6. (**A**) Relative percentages of rare mutation types determined by SSCS and DCS analyses. (**B-E**) Rare mutation context spectra determined by SSCS and DCS analyses.
