## Supplementary material for "Detection of low-frequency mutations and removal of heat-induced artifactual mutations using Duplex Sequencing"

**Table S6.** Heat-induced artifactual rare mutations identified in the thirteen protein-coding mtDNA genes o

| Gene | DNA mutation | AA Change |
|---|---|---|
| MT-ND1 | C3311A | *Syn* |
| MT-ND1 | C3311T | *Syn* |
| MT-ND1 | C3320G | N5K |
| MT-ND1 | C3321A | L6I |
| MT-ND1 | T3322C | L6P |
| MT-ND1 | C3323A | *Syn* |
| MT-ND1 | T3334A | I10N |
| MT-ND1 | A3338G | *Syn* |
| MT-ND1 | C3340A | P12H |
| MT-ND1 | G3356A | *Syn* |
| MT-ND1 | G3356T | M17I |
| MT-ND1 | G3357A | A18T |
| MT-ND1 | G3357T | A18S |
| MT-ND1 | C3362A | F19L |
| MT-ND1 | T3364C | L20P |
| MT-ND1 | A3366G | M21V |
| MT-ND1 | C3369A | L22I |
| MT-ND1 | C3374A | *Syn* |
| MT-ND1 | C3374T | *Syn* |
| MT-ND1 | T3385A | I27N |
| MT-ND1 | C3387A | L28M |
| MT-ND1 | G3390T | G29C |
| MT-ND1 | T3395A | Y30Ter |
| MT-ND1 | T3395C | *Syn* |
| MT-ND1 | C3407A | *Syn* |
| MT-ND1 | A3408T | K35Ter |
| MT-ND1 | G3411A | G36S |
| MT-ND1 | T3424A | V40E |
| MT-ND1 | T3424C | V40A |
| MT-ND1 | G3426A | G41S |
| MT-ND1 | G3426C | G41R |
| MT-ND1 | G3426T | G41C |
| MT-ND1 | C3431A | *Syn* |
| MT-ND1 | C3431T | *Syn* |
| MT-ND1 | G3436A | G44E |
| MT-ND1 | G3436T | G44V |
| MT-ND1 | C3438A | L45M |
| MT-ND1 | C3444A | Q47K |
| MT-ND1 | C3452A | F49L |
| MT-ND1 | G3453A | A50T |
| MT-ND1 | G3453C | A50P |
| MT-ND1 | G3453T | A50S |
| MT-ND1 | C3460A | A52D |
| MT-ND1 | C3460T | A52V |
| MT-ND1 | C3461A | *Syn* |
| MT-ND1 | C3461G | *Syn* |
| MT-ND1 | G3480A | E59K |
| MT-ND1 | G3480T | E59Ter |
| MT-ND1 | C3483A | P60T |
| MT-ND1 | C3484A | P60H |
| MT-ND1 | C3486A | L61M |

| | | |
|---|---|---|
| MT-ND1 | C3492T | P63S |
| MT-ND1 | T3501A | S66T |
| MT-ND1 | C3502A | S66Y |
| MT-ND1 | C3505T | T67I |
| MT-ND1 | C3513A | L70I |
| MT-ND1 | C3523T | T73I |
| MT-ND1 | C3524A | *Syn* |
| MT-ND1 | C3524T | *Syn* |
| MT-ND1 | C3526A | A74D |
| MT-ND1 | C3527A | *Syn* |
| MT-ND1 | C3540A | L79I |
| MT-ND1 | C3540T | L79F |
| MT-ND1 | C3545A | *Syn* |
| MT-ND1 | C3548A | I81M |
| MT-ND1 | C3550A | A82D |
| MT-ND1 | G3562T | W86L |
| MT-ND1 | C3566A | *Syn* |
| MT-ND1 | T3571C | L89P |
| MT-ND1 | C3574A | P90H |
| MT-ND1 | A3582C | N93H |
| MT-ND1 | C3585T | P94S |
| MT-ND1 | C3587T | *Syn* |
| MT-ND1 | C3588A | L95M |
| MT-ND1 | T3589C | L95P |
| MT-ND1 | C3611T | *Syn* |
| MT-ND1 | T3619C | I105T |
| MT-ND1 | A3627C | T108P |
| MT-ND1 | C3629A | *Syn* |
| MT-ND1 | T3630C | S109P |
| MT-ND1 | C3635A | S110Ter |
| MT-ND1 | C3635T | *Syn* |
| MT-ND1 | C3636A | L111M |
| MT-ND1 | C3640A | A112D |
| MT-ND1 | T3655A | L117H |
| MT-ND1 | G3658A | W118Ter |
| MT-ND1 | G3658C | W118S |
| MT-ND1 | G3658T | W118L |
| MT-ND1 | C3661A | S119Ter |
| MT-ND1 | G3663T | G120W |
| MT-ND1 | C3670A | A122E |
| MT-ND1 | C3670T | A122V |
| MT-ND1 | A3685C | Y127S |
| MT-ND1 | C3689A | *Syn* |
| MT-ND1 | T3694C | I130T |
| MT-ND1 | C3695A | I130M |
| MT-ND1 | G3697A | G131D |
| MT-ND1 | G3697T | G131V |
| MT-ND1 | C3698A | *Syn* |
| MT-ND1 | A3701T | *Syn* |
| MT-ND1 | G3704T | *Syn* |
| MT-ND1 | C3705A | R134Ter |
| MT-ND1 | G3708T | A135S |
| MT-ND1 | A3710T | *Syn* |
| MT-ND1 | T3712C | V136A |
| MT-ND1 | G3714A | A137T |

| | | |
|---|---|---|
| MT-ND1 | T3726C | S141P |
| MT-ND1 | C3727A | S141Ter |
| MT-ND1 | C3737A | *Syn* |
| MT-ND1 | C3745A | A147D |
| MT-ND1 | C3746A | *Syn* |
| MT-ND1 | C3749A | I148M |
| MT-ND1 | C3753A | L150M |
| MT-ND1 | T3790C | L162P |
| MT-ND1 | C3793A | S163Y |
| MT-ND1 | C3793T | S163F |
| MT-ND1 | C3797A | *Syn* |
| MT-ND1 | C3819A | L172I |
| MT-ND1 | T3829C | L175P |
| MT-ND1 | C3830A | *Syn* |
| MT-ND1 | C3830G | *Syn* |
| MT-ND1 | C3835A | P177Q |
| MT-ND1 | C3838A | S178Ter |
| MT-ND1 | A3842C | W179C |
| MT-ND1 | C3843T | P180S |
| MT-ND1 | C3844A | P180H |
| MT-ND1 | G3848T | L181F |
| MT-ND1 | G3849T | A182S |
| MT-ND1 | C3850A | A182D |
| MT-ND1 | A3860T | W185C |
| MT-ND1 | C3877A | A191E |
| MT-ND1 | G3881T | E192D |
| MT-ND1 | C3950A | Y215Ter |
| MT-ND1 | C3950T | *Syn* |
| MT-ND1 | C3952A | A216D |
| MT-ND1 | C3953G | *Syn* |
| MT-ND1 | G3954A | A217T |
| MT-ND1 | G3954T | A217S |
| MT-ND1 | T3963C | F220L |
| MT-ND1 | C3965A | F220L |
| MT-ND1 | C3974A | F223L |
| MT-ND1 | C3974T | *Syn* |
| MT-ND1 | C3977A | F224L |
| MT-ND1 | C3983A | *Syn* |
| MT-ND1 | G3984T | E227Ter |
| MT-ND1 | C3989T | *Syn* |
| MT-ND1 | C4049A | D248E |
| MT-ND1 | C4051A | A249E |
| MT-ND1 | T4056C | S251P |
| MT-ND1 | C4058A | *Syn* |
| MT-ND1 | C4085A | *Syn* |
| MT-ND1 | C4088A | *Syn* |
| MT-ND1 | C4093T | T263I |
| MT-ND1 | C4094A | *Syn* |
| MT-ND1 | C4095A | L264M |
| MT-ND1 | T4102C | L266P |
| MT-ND1 | C4105T | T267I |
| MT-ND1 | C4109A | *Syn* |
| MT-ND1 | C4151T | *Syn* |
| MT-ND1 | T4159C | L285P |
| MT-ND1 | C4167A | L288I |

| | | |
|---|---|---|
| MT-ND1 | G4174T | W290L |
| MT-ND1 | C4184A | F293L |
| MT-ND1 | C4185T | *Syn* |
| MT-ND1 | C4189A | P295Q |
| MT-ND1 | C4196A | *Syn* |
| MT-ND1 | C4196T | *Syn* |
| MT-ND1 | T4198C | L298P |
| MT-ND2 | T4474A | N2K |
| MT-ND2 | T4474C | *Syn* |
| MT-ND2 | G4480A | *Syn* |
| MT-ND2 | G4480T | *Syn* |
| MT-ND2 | C4512T | A15V |
| MT-ND2 | G4514T | G16C |
| MT-ND2 | G4515A | G16D |
| MT-ND2 | T4521C | L18P |
| MT-ND2 | C4527T | T20M |
| MT-ND2 | G4529T | A21S |
| MT-ND2 | G4531A | *Syn* |
| MT-ND2 | G4531T | *Syn* |
| MT-ND2 | C4543T | *Syn* |
| MT-ND2 | G4545T | W26L |
| MT-ND2 | A4553T | T29S |
| MT-ND2 | C4554A | T29N |
| MT-ND2 | G4557T | W30L |
| MT-ND2 | G4559A | V31M |
| MT-ND2 | C4564T | *Syn* |
| MT-ND2 | G4568A | E34K |
| MT-ND2 | T4578A | M37K |
| MT-ND2 | G4579A | *Syn* |
| MT-ND2 | G4579T | M37I |
| MT-ND2 | C4592A | P42T |
| MT-ND2 | C4602A | T45N |
| MT-ND2 | C4603A | *Syn* |
| MT-ND2 | T4611A | M48K |
| MT-ND2 | C4617A | P50H |
| MT-ND2 | T4622C | S52P |
| MT-ND2 | G4631A | A55T |
| MT-ND2 | G4631T | A55S |
| MT-ND2 | C4635A | A56D |
| MT-ND2 | G4654A | *Syn* |
| MT-ND2 | G4654T | *Syn* |
| MT-ND2 | C4662T | T65I |
| MT-ND2 | C4675A | I69M |
| MT-ND2 | A4682G | M72V |
| MT-ND2 | G4685T | A73S |
| MT-ND2 | C4690A | I74M |
| MT-ND2 | C4690T | *Syn* |
| MT-ND2 | C4691A | L75I |
| MT-ND2 | C4708A | *Syn* |
| MT-ND2 | C4710A | S81Y |
| MT-ND2 | C4715A | Q83K |
| MT-ND2 | A4716T | Q83L |
| MT-ND2 | C4723A | *Syn* |
| MT-ND2 | C4728A | T87N |
| MT-ND2 | T4735A | *Syn* |

| | | |
|---|---|---|
| MT-ND2 | C4737A | T90N |
| MT-ND2 | C4738A | *Syn* |
| MT-ND2 | C4738G | *Syn* |
| MT-ND2 | A4739T | N91Y |
| MT-ND2 | C4752A | S95Ter |
| MT-ND2 | A4753T | *Syn* |
| MT-ND2 | C4762A | I98M |
| MT-ND2 | T4767A | M100K |
| MT-ND2 | C4770A | A101D |
| MT-ND2 | G4787A | G107Ter |
| MT-ND2 | G4793T | A109S |
| MT-ND2 | C4798A | *Syn* |
| MT-ND2 | C4807T | *Syn* |
| MT-ND2 | T4812C | V115A |
| MT-ND2 | C4813A | *Syn* |
| MT-ND2 | C4815A | P116Q |
| MT-ND2 | G4817A | E117K |
| MT-ND2 | A4818G | E117G |
| MT-ND2 | G4820A | V118I |
| MT-ND2 | C4834A | *Syn* |
| MT-ND2 | C4834T | *Syn* |
| MT-ND2 | C4836A | P123H |
| MT-ND2 | G4840T | *Syn* |
| MT-ND2 | C4845A | S126Y |
| MT-ND2 | C4846A | *Syn* |
| MT-ND2 | C4846T | *Syn* |
| MT-ND2 | C4846G | *Syn* |
| MT-ND2 | G4847T | G127C |
| MT-ND2 | C4850A | L128M |
| MT-ND2 | C4850T | *Syn* |
| MT-ND2 | G4852T | *Syn* |
| MT-ND2 | C4853T | L129F |
| MT-ND2 | T4865C | W133R |
| MT-ND2 | G4866A | W133Ter |
| MT-ND2 | G4866C | W133S |
| MT-ND2 | C4878A | A137D |
| MT-ND2 | C4887A | S140Ter |
| MT-ND2 | C4905A | S146Y |
| MT-ND2 | C4907A | P147T |
| MT-ND2 | A4921C | *Syn* |
| MT-ND2 | C4925A | L153I |
| MT-ND2 | C4935T | T156I |
| MT-ND2 | C4945A | I159M |
| MT-ND2 | T4947C | L160S |
| MT-ND2 | C4950A | S161Y |
| MT-ND2 | C4951A | *Syn* |
| MT-ND2 | C4951T | *Syn* |
| MT-ND2 | C4954A | I162M |
| MT-ND2 | G4968A | W167Ter |
| MT-ND2 | G4968T | W167L |
| MT-ND2 | G4971A | G168D |
| MT-ND2 | G4971T | G168V |
| MT-ND2 | G4973A | G169Ter |
| MT-ND2 | G4973T | G169W |
| MT-ND2 | G4974A | G169E |

| | | |
|---|---|---|
| MT-ND2 | G4974C | G169A |
| MT-ND2 | C4981T | *Syn* |
| MT-ND2 | C4982A | Q172K |
| MT-ND2 | C4982T | Q172Ter |
| MT-ND2 | C4988A | Q174K |
| MT-ND2 | G4990T | Q174H |
| MT-ND2 | A5005T | L179F |
| MT-ND2 | C5016A | S183Ter |
| MT-ND2 | A5021C | T185P |
| MT-ND2 | C5022T | T185I |
| MT-ND2 | A5025G | H186R |
| MT-ND2 | C5026A | H186Q |
| MT-ND2 | T5033A | W189Ter |
| MT-ND2 | G5034C | W189S |
| MT-ND2 | G5034T | W189L |
| MT-ND2 | G5042A | A192T |
| MT-ND2 | T5047C | *Syn* |
| MT-ND2 | C5051A | P195T |
| MT-ND2 | C5051G | P195A |
| MT-ND2 | A5058C | N197T |
| MT-ND2 | C5065A | N199K |
| MT-ND2 | C5085T | T206I |
| MT-ND2 | A5087T | I207F |
| MT-ND2 | C5098T | *Syn* |
| MT-ND2 | C5099A | L211M |
| MT-ND2 | C5110A | *Syn* |
| MT-ND2 | C5110T | *Syn* |
| MT-ND2 | C5137G | *Syn* |
| MT-ND2 | C5193A | P242H |
| MT-ND2 | C5201T | P245S |
| MT-ND2 | C5202A | P245Q |
| MT-ND2 | T5204C | S246P |
| MT-ND2 | C5205A | S246Y |
| MT-ND2 | C5213A | L249I |
| MT-ND2 | C5213T | L249F |
| MT-ND2 | C5213G | L249V |
| MT-ND2 | C5215A | *Syn* |
| MT-ND2 | C5217A | S250Y |
| MT-ND2 | T5220C | L251P |
| MT-ND2 | G5226A | G253D |
| MT-ND2 | G5226T | G253V |
| MT-ND2 | C5235A | P256Q |
| MT-ND2 | C5235T | P256L |
| MT-ND2 | C5235G | P256R |
| MT-ND2 | C5242A | *Syn* |
| MT-ND2 | C5245A | *Syn* |
| MT-ND2 | G5251T | L261F |
| MT-ND2 | C5252A | P262T |
| MT-ND2 | C5252T | P262S |
| MT-ND2 | G5260T | W264C |
| MT-ND2 | C5262A | A265D |
| MT-ND2 | C5262T | A265V |
| MT-ND2 | T5277A | F270Y |
| MT-ND2 | C5278A | F270L |
| MT-ND2 | C5278T | *Syn* |

| | | |
|---|---|---|
| MT-ND2 | C5280T | T271M |
| MT-ND2 | C5287A | N273K |
| MT-ND2 | G5292C | S275T |
| MT-ND2 | G5292T | S275I |
| MT-ND2 | C5293A | S275Ter |
| MT-ND2 | C5293T | *Syn* |
| MT-ND2 | C5296A | *Syn* |
| MT-ND2 | C5296G | *Syn* |
| MT-ND2 | C5299A | I277M |
| MT-ND2 | C5299T | *Syn* |
| MT-ND2 | C5302A | I278M |
| MT-ND2 | C5302T | *Syn* |
| MT-ND2 | C5303A | P279T |
| MT-ND2 | C5307A | T280N |
| MT-ND2 | C5307T | T280I |
| MT-ND2 | T5310A | I281N |
| MT-ND2 | C5311A | I281M |
| MT-ND2 | C5316A | A283D |
| MT-ND2 | A5321G | I285V |
| MT-ND2 | C5323A | I285M |
| MT-ND2 | T5331A | L288H |
| MT-ND2 | T5331C | L288P |
| MT-ND2 | C5338A | *Syn* |
| MT-ND2 | T5345C | Y293H |
| MT-ND2 | C5348T | *Syn* |
| MT-ND2 | T5382C | L305P |
| MT-ND2 | C5391A | S308Y |
| MT-ND2 | A5431C | K321N |
| MT-ND2 | C5433A | P322H |
| MT-ND2 | C5439A | P324Q |
| MT-ND2 | A5440T | *Syn* |
| MT-ND2 | C5443A | F325L |
| MT-ND2 | T5445C | L326P |
| MT-ND2 | C5453A | L329I |
| MT-ND2 | T5454A | L329H |
| MT-ND2 | C5458A | I330M |
| MT-ND2 | C5458T | *Syn* |
| MT-ND2 | C5458G | I330M |
| MT-ND2 | C5460A | A331D |
| MT-ND2 | C5460T | A331V |
| MT-ND2 | C5461A | *Syn* |
| MT-ND2 | C5462A | L332I |
| MT-ND2 | C5462T | L332F |
| MT-ND2 | G5470T | *Syn* |
| MT-ND2 | C5471A | L335M |
| MT-ND2 | C5477A | L337M |
| MT-ND2 | T5486C | S340P |
| MT-ND2 | C5488T | *Syn* |
| MT-ND2 | C5490G | P341R |
| MT-ND2 | T5491C | *Syn* |
| MT-ND2 | T5505C | I346T |
| MT-CO1 | C5926T | *Syn* |
| MT-CO1 | T5946C | I15T |
| MT-CO1 | C5952A | T17K |
| MT-CO1 | A5958C | Y19S |

| | | |
|---|---|---|
| MT-CO1 | C5960A | L20M |
| MT-CO1 | C5960T | *Syn* |
| MT-CO1 | C5973A | A24E |
| MT-CO1 | G5976A | W25Ter |
| MT-CO1 | G5976T | W25L |
| MT-CO1 | C5986A | *Syn* |
| MT-CO1 | T5998C | *Syn* |
| MT-CO1 | C5999T | *Syn* |
| MT-CO1 | C6005A | L35I |
| MT-CO1 | C6005T | L35F |
| MT-CO1 | C6019A | *Syn* |
| MT-CO1 | G6022C | E40D |
| MT-CO1 | G6022T | E40D |
| MT-CO1 | C6023A | L41M |
| MT-CO1 | C6023T | *Syn* |
| MT-CO1 | G6025C | *Syn* |
| MT-CO1 | G6025T | *Syn* |
| MT-CO1 | C6028A | *Syn* |
| MT-CO1 | C6029A | Q43K |
| MT-CO1 | C6029G | Q43E |
| MT-CO1 | C6032A | P44T |
| MT-CO1 | C6033A | P44Q |
| MT-CO1 | G6036A | G45D |
| MT-CO1 | G6036T | G45V |
| MT-CO1 | G6047T | G49C |
| MT-CO1 | G6048A | G49D |
| MT-CO1 | T6049G | *Syn* |
| MT-CO1 | G6053A | D51N |
| MT-CO1 | G6053C | D51H |
| MT-CO1 | G6053T | D51Y |
| MT-CO1 | C6055A | D51E |
| MT-CO1 | C6058A | H52Q |
| MT-CO1 | T6060C | I53T |
| MT-CO1 | C6064A | Y54Ter |
| MT-CO1 | T6075A | V58D |
| MT-CO1 | C6082A | *Syn* |
| MT-CO1 | C6082T | *Syn* |
| MT-CO1 | A6084C | H61P |
| MT-CO1 | T6085A | H61Q |
| MT-CO1 | C6133T | *Syn* |
| MT-CO1 | G6167T | A89S |
| MT-CO1 | C6170A | P90T |
| MT-CO1 | A6174T | D91V |
| MT-CO1 | T6177A | M92K |
| MT-CO1 | G6179T | A93S |
| MT-CO1 | T6184C | *Syn* |
| MT-CO1 | A6194G | N98D |
| MT-CO1 | C6212A | L104I |
| MT-CO1 | C6212T | L104F |
| MT-CO1 | T6213C | L104P |
| MT-CO1 | T6220C | *Syn* |
| MT-CO1 | C6221A | P107T |
| MT-CO1 | C6222A | P107H |
| MT-CO1 | C6222T | P107L |
| MT-CO1 | C6222G | P107R |

| | | |
|---|---|---|
| MT-CO1 | C6223A | *Syn* |
| MT-CO1 | C6223T | *Syn* |
| MT-CO1 | C6227A | L109I |
| MT-CO1 | G6238C | *Syn* |
| MT-CO1 | G6238T | *Syn* |
| MT-CO1 | C6241A | *Syn* |
| MT-CO1 | C6241T | *Syn* |
| MT-CO1 | G6242T | A114S |
| MT-CO1 | C6246A | S115Y |
| MT-CO1 | G6260A | A120T |
| MT-CO1 | G6260T | A120S |
| MT-CO1 | G6276A | G125D |
| MT-CO1 | G6276T | G125V |
| MT-CO1 | C6286A | *Syn* |
| MT-CO1 | C6286T | *Syn* |
| MT-CO1 | C6289T | *Syn* |
| MT-CO1 | C6290A | P130T |
| MT-CO1 | C6291A | P130H |
| MT-CO1 | T6296C | *Syn* |
| MT-CO1 | G6303T | G134V |
| MT-CO1 | C6307T | *Syn* |
| MT-CO1 | C6312A | S137Y |
| MT-CO1 | C6313A | *Syn* |
| MT-CO1 | A6315C | H138P |
| MT-CO1 | C6317A | P139T |
| MT-CO1 | G6321T | G140V |
| MT-CO1 | C6328A | *Syn* |
| MT-CO1 | C6328G | *Syn* |
| MT-CO1 | C6334T | *Syn* |
| MT-CO1 | C6339A | T146N |
| MT-CO1 | C6343A | I147M |
| MT-CO1 | C6349G | *Syn* |
| MT-CO1 | C6355A | H151Q |
| MT-CO1 | C6356A | L152M |
| MT-CO1 | G6362A | G154S |
| MT-CO1 | G6362T | G154C |
| MT-CO1 | C6367A | *Syn* |
| MT-CO1 | C6369A | S156Y |
| MT-CO1 | C6385A | *Syn* |
| MT-CO1 | A6398T | T166S |
| MT-CO1 | C6399A | T166K |
| MT-CO1 | A6401T | T167S |
| MT-CO1 | T6408A | I169N |
| MT-CO1 | C6409A | I169M |
| MT-CO1 | T6414A | M171K |
| MT-CO1 | C6419G | P173A |
| MT-CO1 | C6423A | P174H |
| MT-CO1 | C6432A | T177N |
| MT-CO1 | C6433A | *Syn* |
| MT-CO1 | C6433G | *Syn* |
| MT-CO1 | C6448A | *Syn* |
| MT-CO1 | C6457A | *Syn* |
| MT-CO1 | C6462A | S187Y |
| MT-CO1 | C6462G | S187C |
| MT-CO1 | C6463A | *Syn* |

| | | |
|---|---|---|
| MT-CO1 | A6478T | *Syn* |
| MT-CO1 | C6488T | L196F |
| MT-CO1 | T6492C | L197P |
| MT-CO1 | T6498C | L199P |
| MT-CO1 | C6500A | P200T |
| MT-CO1 | C6500G | P200A |
| MT-CO1 | C6506A | L202M |
| MT-CO1 | C6506T | *Syn* |
| MT-CO1 | C6510A | A203D |
| MT-CO1 | G6515A | G205S |
| MT-CO1 | G6515C | G205R |
| MT-CO1 | G6515T | G205C |
| MT-CO1 | T6519A | I206N |
| MT-CO1 | C6522A | T207N |
| MT-CO1 | C6522T | T207I |
| MT-CO1 | C6527T | *Syn* |
| MT-CO1 | C6530A | L210M |
| MT-CO1 | C6530T | *Syn* |
| MT-CO1 | G6536C | D212H |
| MT-CO1 | G6536T | D212Y |
| MT-CO1 | G6540A | R213H |
| MT-CO1 | G6540T | R213L |
| MT-CO1 | C6550A | N216K |
| MT-CO1 | C6550T | *Syn* |
| MT-CO1 | C6552A | T217N |
| MT-CO1 | C6555T | T218I |
| MT-CO1 | C6556A | *Syn* |
| MT-CO1 | C6556T | *Syn* |
| MT-CO1 | C6556G | *Syn* |
| MT-CO1 | C6568A | *Syn* |
| MT-CO1 | C6570A | A223D |
| MT-CO1 | C6570T | A223V |
| MT-CO1 | G6576T | G225V |
| MT-CO1 | G6578T | G226W |
| MT-CO1 | T6591C | L230P |
| MT-CO1 | T6605A | F235I |
| MT-CO1 | C6622T | *Syn* |
| MT-CO1 | T6625C | *Syn* |
| MT-CO1 | G6629T | V243F |
| MT-CO1 | T6630C | V243A |
| MT-CO1 | C6638A | L246I |
| MT-CO1 | G6650T | G250C |
| MT-CO1 | T6672A | I257N |
| MT-CO1 | C6688A | *Syn* |
| MT-CO1 | G6721T | M273I |
| MT-CO1 | G6722T | V274F |
| MT-CO1 | C6724A | *Syn* |
| MT-CO1 | A6731T | M277L |
| MT-CO1 | T6732C | M277T |
| MT-CO1 | C6738A | S279Ter |
| MT-CO1 | T6741C | I280T |
| MT-CO1 | G6743A | G281S |
| MT-CO1 | G6743C | G281R |
| MT-CO1 | G6743T | G281C |
| MT-CO1 | G6744T | G281V |

| | | |
|---|---|---|
| MT-CO1 | G6752T | G284W |
| MT-CO1 | C6760A | I286M |
| MT-CO1 | G6761A | V287M |
| MT-CO1 | G6765T | W288L |
| MT-CO1 | A6766T | W288C |
| MT-CO1 | G6767A | A289T |
| MT-CO1 | G6785A | V295M |
| MT-CO1 | G6785T | V295L |
| MT-CO1 | G6789T | G296V |
| MT-CO1 | C6796A | D298E |
| MT-CO1 | G6797C | V299L |
| MT-CO1 | G6797T | V299L |
| MT-CO1 | C6806G | R302G |
| MT-CO1 | G6807A | R302Q |
| MT-CO1 | G6807T | R302L |
| MT-CO1 | C6819A | T306N |
| MT-CO1 | C6820A | *Syn* |
| MT-CO1 | C6822A | S307Y |
| MT-CO1 | C6823A | *Syn* |
| MT-CO1 | T6826A | *Syn* |
| MT-CO1 | C6829A | *Syn* |
| MT-CO1 | C6838A | I312M |
| MT-CO1 | G6839C | A313P |
| MT-CO1 | G6839T | A313S |
| MT-CO1 | C6847A | *Syn* |
| MT-CO1 | C6847T | *Syn* |
| MT-CO1 | A6848C | T316P |
| MT-CO1 | C6850A | *Syn* |
| MT-CO1 | G6851T | G317C |
| MT-CO1 | G6852T | G317V |
| MT-CO1 | A6871G | *Syn* |
| MT-CO1 | C6877A | *Syn* |
| MT-CO1 | C6877G | *Syn* |
| MT-CO1 | A6880C | *Syn* |
| MT-CO1 | T6882C | L327P |
| MT-CO1 | C6892T | *Syn* |
| MT-CO1 | C6906A | S335Y |
| MT-CO1 | C6906T | S335F |
| MT-CO1 | T6927C | L342P |
| MT-CO1 | A6931G | *Syn* |
| MT-CO1 | C6934A | F344L |
| MT-CO1 | C6937A | I345M |
| MT-CO1 | C6949A | *Syn* |
| MT-CO1 | C6949T | *Syn* |
| MT-CO1 | G6950T | V350L |
| MT-CO1 | G6957T | G352V |
| MT-CO1 | G6966T | G355V |
| MT-CO1 | C6967T | *Syn* |
| MT-CO1 | G6977T | A359S |
| MT-CO1 | C6978A | A359E |
| MT-CO1 | C6984A | S361Ter |
| MT-CO1 | C6989T | *Syn* |
| MT-CO1 | G6992C | D364H |
| MT-CO1 | C7001T | *Syn* |
| MT-CO1 | C7006T | *Syn* |

| | | |
|---|---|---|
| MT-CO1 | C7009T | *Syn* |
| MT-CO1 | G7025T | A375S |
| MT-CO1 | C7033G | F377L |
| MT-CO1 | C7034A | H378N |
| MT-CO1 | C7043T | *Syn* |
| MT-CO1 | C7047A | S382Ter |
| MT-CO1 | C7047T | S382L |
| MT-CO1 | G7052A | G384Ter |
| MT-CO1 | C7056A | A385D |
| MT-CO1 | C7056T | A385V |
| MT-CO1 | T7057C | *Syn* |
| MT-CO1 | C7065A | A388D |
| MT-CO1 | C7066A | *Syn* |
| MT-CO1 | C7066T | *Syn* |
| MT-CO1 | G7076T | G392C |
| MT-CO1 | G7077A | G392D |
| MT-CO1 | C7081T | *Syn* |
| MT-CO1 | A7090T | W396C |
| MT-CO1 | T7093C | *Syn* |
| MT-CO1 | C7095A | P398H |
| MT-CO1 | C7096T | *Syn* |
| MT-CO1 | C7102T | *Syn* |
| MT-CO1 | A7105G | *Syn* |
| MT-CO1 | G7106A | G402S |
| MT-CO1 | G7106C | G402R |
| MT-CO1 | G7106T | G402C |
| MT-CO1 | C7120T | *Syn* |
| MT-CO1 | C7121A | Q407K |
| MT-CO1 | A7124C | T408P |
| MT-CO1 | C7125A | T408N |
| MT-CO1 | C7126A | *Syn* |
| MT-CO1 | A7128G | Y409C |
| MT-CO1 | C7129A | Y409Ter |
| MT-CO1 | C7129T | *Syn* |
| MT-CO1 | C7131A | A410D |
| MT-CO1 | C7139A | H413N |
| MT-CO1 | C7139T | H413Y |
| MT-CO1 | A7140T | H413L |
| MT-CO1 | C7144A | F414L |
| MT-CO1 | G7163T | V421L |
| MT-CO1 | A7183G | *Syn* |
| MT-CO1 | C7187A | H429N |
| MT-CO1 | C7187T | H429Y |
| MT-CO1 | G7197A | G432D |
| MT-CO1 | C7203A | S434Y |
| MT-CO1 | C7214A | R438Ter |
| MT-CO1 | G7215T | R438L |
| MT-CO1 | C7222A | Y440Ter |
| MT-CO1 | G7225T | *Syn* |
| MT-CO1 | G7226C | D442H |
| MT-CO1 | G7226T | D442Y |
| MT-CO1 | C7232A | P444T |
| MT-CO1 | C7246A | *Syn* |
| MT-CO1 | C7259A | L453M |
| MT-CO1 | C7259T | *Syn* |

| | | |
|---|---|---|
| MT-CO1 | T7260C | L453P |
| MT-CO1 | C7266A | S455Y |
| MT-CO1 | C7273A | *Syn* |
| MT-CO1 | C7275A | S458Ter |
| MT-CO1 | C7275T | S458L |
| MT-CO1 | A7276T | *Syn* |
| MT-CO1 | A7280T | I460F |
| MT-CO1 | C7290A | T463K |
| MT-CO1 | C7290T | T463M |
| MT-CO1 | G7292T | A464S |
| MT-CO1 | C7293A | A464E |
| MT-CO1 | G7295C | V465L |
| MT-CO1 | G7295T | V465L |
| MT-CO1 | T7311C | F470S |
| MT-CO1 | C7312A | F470L |
| MT-CO1 | G7315T | M471I |
| MT-CO1 | C7326A | A475D |
| MT-CO1 | C7326T | A475V |
| MT-CO1 | C7327A | *Syn* |
| MT-CO1 | C7327T | *Syn* |
| MT-CO2 | G7594T | A4S |
| MT-CO2 | C7595A | A4E |
| MT-CO2 | C7598A | A5E |
| MT-CO2 | C7609A | L9M |
| MT-CO2 | C7612A | Q10K |
| MT-CO2 | C7622T | T13I |
| MT-CO2 | T7623C | *Syn* |
| MT-CO2 | C7632A | I16M |
| MT-CO2 | C7654A | H24N |
| MT-CO2 | C7660A | H26N |
| MT-CO2 | T7667C | L28P |
| MT-CO2 | C7668A | *Syn* |
| MT-CO2 | C7668G | *Syn* |
| MT-CO2 | C7681T | L33F |
| MT-CO2 | T7685A | I34N |
| MT-CO2 | C7686A | I34M |
| MT-CO2 | G7688A | C35Y |
| MT-CO2 | G7688T | C35F |
| MT-CO2 | C7693A | L37M |
| MT-CO2 | C7693G | L37V |
| MT-CO2 | G7696A | V38I |
| MT-CO2 | C7698A | *Syn* |
| MT-CO2 | G7701A | *Syn* |
| MT-CO2 | T7704A | Y40Ter |
| MT-CO2 | C7706T | A41V |
| MT-CO2 | C7714A | L44M |
| MT-CO2 | C7714T | *Syn* |
| MT-CO2 | C7720T | L46F |
| MT-CO2 | C7724T | T47M |
| MT-CO2 | T7740A | N52K |
| MT-CO2 | C7746A | N54K |
| MT-CO2 | C7749A | I55M |
| MT-CO2 | C7749T | *Syn* |
| MT-CO2 | C7751A | S56Ter |
| MT-CO2 | G7753A | D57N |

| | | |
|---|---|---|
| MT-CO2 | G7756A | A58T |
| MT-CO2 | G7756T | A58S |
| MT-CO2 | C7759A | Q59K |
| MT-CO2 | C7786A | L68M |
| MT-CO2 | A7795C | I71L |
| MT-CO2 | T7796C | I71T |
| MT-CO2 | C7797A | I71M |
| MT-CO2 | C7800A | I72M |
| MT-CO2 | C7806A | *Syn* |
| MT-CO2 | T7808A | L75H |
| MT-CO2 | T7808C | L75P |
| MT-CO2 | C7809A | *Syn* |
| MT-CO2 | T7817C | L78P |
| MT-CO2 | C7820A | P79Q |
| MT-CO2 | C7820G | P79R |
| MT-CO2 | C7824A | *Syn* |
| MT-CO2 | T7826C | L81P |
| MT-CO2 | G7829A | R82H |
| MT-CO2 | G7829T | R82L |
| MT-CO2 | T7841A | M86K |
| MT-CO2 | C7844A | T87K |
| MT-CO2 | G7851A | *Syn* |
| MT-CO2 | G7851T | E89D |
| MT-CO2 | C7854A | *Syn* |
| MT-CO2 | C7857T | *Syn* |
| MT-CO2 | C7861A | P93T |
| MT-CO2 | T7864C | S94P |
| MT-CO2 | C7865A | S94Y |
| MT-CO2 | C7880A | S99Ter |
| MT-CO2 | C7880T | S99L |
| MT-CO2 | C7890T | *Syn* |
| MT-CO2 | G7901T | W106L |
| MT-CO2 | C7914A | Y110Ter |
| MT-CO2 | C7917A | *Syn* |
| MT-CO2 | C7923T | *Syn* |
| MT-CO2 | C7926A | *Syn* |
| MT-CO2 | G7928T | G115V |
| MT-CO2 | C7930A | L116M |
| MT-CO2 | T7931C | L116P |
| MT-CO2 | T7934C | I117T |
| MT-CO2 | A7939G | N119D |
| MT-CO2 | T7942C | S120P |
| MT-CO2 | C7943A | S120Y |
| MT-CO2 | T7945C | Y121H |
| MT-CO2 | A7950C | M122I |
| MT-CO2 | T7953C | *Syn* |
| MT-CO2 | C7955A | P124H |
| MT-CO2 | C7956A | *Syn* |
| MT-CO2 | T7960C | *Syn* |
| MT-CO2 | T7964C | F127S |
| MT-CO2 | C7966A | L128M |
| MT-CO2 | G7969T | E129Ter |
| MT-CO2 | A7971G | *Syn* |
| MT-CO2 | C7972A | P130T |
| MT-CO2 | G7976A | G131D |

| | | |
|---|---|---|
| MT-CO2 | A7979C | D132A |
| MT-CO2 | G7985A | R134Q |
| MT-CO2 | G7985T | R134L |
| MT-CO2 | C7987G | L135V |
| MT-CO2 | C7989T | *Syn* |
| MT-CO2 | C7990A | L136I |
| MT-CO2 | G7993T | D137Y |
| MT-CO2 | C7995T | *Syn* |
| MT-CO2 | T7997C | V138A |
| MT-CO2 | A8000T | D139V |
| MT-CO2 | G8011A | V143M |
| MT-CO2 | T8015A | L144H |
| MT-CO2 | T8021A | I146N |
| MT-CO2 | G8023C | E147Q |
| MT-CO2 | G8023T | E147Ter |
| MT-CO2 | C8028A | *Syn* |
| MT-CO2 | C8051A | S156Ter |
| MT-CO2 | A8055T | Q157H |
| MT-CO2 | C8058A | D158E |
| MT-CO2 | C8058T | *Syn* |
| MT-CO2 | C8065A | H161N |
| MT-CO2 | C8065T | H161Y |
| MT-CO2 | G8072T | W163L |
| MT-CO2 | C8079A | *Syn* |
| MT-CO2 | A8083C | T167P |
| MT-CO2 | C8111A | P176H |
| MT-CO2 | C8125A | Q181K |
| MT-CO2 | C8130A | *Syn* |
| MT-CO2 | C8130T | *Syn* |
| MT-CO2 | A8131C | T183P |
| MT-CO2 | C8139A | *Syn* |
| MT-CO2 | C8139T | *Syn* |
| MT-CO2 | C8146T | R188W |
| MT-CO2 | C8150A | P189Q |
| MT-CO2 | A8159G | Y192C |
| MT-CO2 | C8167A | Q195K |
| MT-CO2 | G8171A | C196Y |
| MT-CO2 | G8171T | C196F |
| MT-CO2 | C8172A | C196W |
| MT-CO2 | C8172T | *Syn* |
| MT-CO2 | G8176A | E198K |
| MT-CO2 | G8176C | E198Q |
| MT-CO2 | G8176T | E198Ter |
| MT-CO2 | C8181A | I199M |
| MT-CO2 | C8181T | *Syn* |
| MT-CO2 | G8188A | A202T |
| MT-CO2 | G8188T | A202S |
| MT-CO2 | C8189A | A202E |
| MT-CO2 | C8189T | A202V |
| MT-CO2 | C8194A | H204N |
| MT-CO2 | G8198A | S205N |
| MT-CO2 | G8198T | S205I |
| MT-CO2 | T8201A | F206Y |
| MT-CO2 | C8207A | P208H |
| MT-CO2 | C8211A | I209M |

| | | |
|---|---|---|
| MT-CO2 | C8211T | *Syn* |
| MT-CO2 | G8218T | E212Ter |
| MT-CO2 | T8226C | *Syn* |
| MT-CO2 | C8227A | P215T |
| MT-CO2 | C8227T | P215S |
| MT-CO2 | C8228A | P215H |
| MT-CO2 | C8230A | L216M |
| MT-CO2 | A8236T | I218F |
| MT-CO2 | C8238A | I218M |
| MT-CO2 | G8249T | G222V |
| MT-CO2 | C8252A | P223H |
| MT-CO2 | C8253A | *Syn* |
| MT-CO2 | A8256T | *Syn* |
| MT-CO2 | C8261T | T226I |
| MT-CO2 | C8262A | *Syn* |
| MT-ATP8 | T8366A | M1K |
| MT-ATP8 | C8369A | P2H |
| MT-ATP8 | C8371A | Q3K |
| MT-ATP8 | G8386A | V8M |
| MT-ATP8 | G8386T | V8L |
| MT-ATP8 | T8389C | W9R |
| MT-ATP8 | G8391T | W9C |
| MT-ATP8 | C8392A | P10T |
| MT-ATP8 | C8393A | P10H |
| MT-ATP8 | C8397A | *Syn* |
| MT-ATP8 | C8397T | *Syn* |
| MT-ATP8 | C8405T | T14I |
| MT-ATP8 | C8406A | *Syn* |
| MT-ATP8 | C8416A | L18I |
| MT-ATP8 | C8416T | L18F |
| MT-ATP8 | C8428A | L22I |
| MT-ATP8 | C8428T | L22F |
| MT-ATP8 | C8433A | I23M |
| MT-ATP8 | C8433T | *Syn* |
| MT-ATP8 | C8435A | T24N |
| MT-ATP8 | T8449C | *Syn* |
| MT-ATP8 | C8454A | N30K |
| MT-ATP8 | C8454T | *Syn* |
| MT-ATP8 | C8463G | Y33Ter |
| MT-ATP8 | C8464A | H34N |
| MT-ATP8 | C8464T | H34Y |
| MT-ATP8 | A8465C | H34P |
| MT-ATP8 | C8467A | L35M |
| MT-ATP8 | C8473A | P37T |
| MT-ATP8 | C8474A | P37H |
| MT-ATP8 | C8475A | *Syn* |
| MT-ATP8 | T8476A | S38T |
| MT-ATP8 | A8483G | K40Ter |
| MT-ATP8 | C8485A | P41T |
| MT-ATP8 | C8486A | P41H |
| MT-ATP8 | C8487T | *Syn* |
| MT-ATP8 | A8490T | M42I |
| MT-ATP8 | C8514A | *Syn* |
| MT-ATP8 | C8514T | *Syn* |
| MT-ATP8 | A8520C | E52D |

| | | |
|---|---|---|
| MT-ATP8 | A8526T | K54N |
| MT-ATP6 | A8526T | M1L |
| MT-ATP8 | C8538A | I58M |
| MT-ATP6 | C8538A | L5M |
| MT-ATP8 | C8557A | P65T |
| MT-ATP6 | C8557A | A11D |
| MT-ATP8 | C8557T | P65S |
| MT-ATP6 | C8557T | A11V |
| MT-ATP8 | C8561A | P66Q |
| MT-ATP6 | C8561A | *Syn* |
| MT-ATP6 | C8574A | L17M |
| MT-ATP6 | C8577A | P18T |
| MT-ATP6 | C8579A | *Syn* |
| MT-ATP6 | C8581A | A19D |
| MT-ATP6 | C8581T | A19V |
| MT-ATP6 | C8582A | *Syn* |
| MT-ATP6 | G8583T | A20S |
| MT-ATP6 | C8598A | L25M |
| MT-ATP6 | C8598G | L25V |
| MT-ATP6 | C8606A | *Syn* |
| MT-ATP6 | C8608A | P28H |
| MT-ATP6 | T8611A | L29Q |
| MT-ATP6 | G8615T | L30F |
| MT-ATP6 | C8620A | P32H |
| MT-ATP6 | C8634A | L37I |
| MT-ATP6 | A8641G | N39S |
| MT-ATP6 | C8645T | *Syn* |
| MT-ATP6 | C8649A | L42M |
| MT-ATP6 | C8654A | I43M |
| MT-ATP6 | C8660A | *Syn* |
| MT-ATP6 | C8679T | *Syn* |
| MT-ATP6 | A8682C | T53P |
| MT-ATP6 | C8708T | *Syn* |
| MT-ATP6 | C8721T | R66W |
| MT-ATP6 | G8722T | R66L |
| MT-ATP6 | A8724C | T67P |
| MT-ATP6 | C8726A | *Syn* |
| MT-ATP6 | C8733A | L70I |
| MT-ATP6 | C8733T | L70F |
| MT-ATP6 | T8745A | S74T |
| MT-ATP6 | A8751T | I76F |
| MT-ATP6 | C8753A | I76M |
| MT-ATP6 | C8764A | A80D |
| MT-ATP6 | C8764T | A80V |
| MT-ATP6 | C8765A | *Syn* |
| MT-ATP6 | C8765T | *Syn* |
| MT-ATP6 | C8767A | T81K |
| MT-ATP6 | C8770A | T82N |
| MT-ATP6 | C8770T | T82I |
| MT-ATP6 | C8778A | L85I |
| MT-ATP6 | C8778T | L85F |
| MT-ATP6 | C8780A | *Syn* |
| MT-ATP6 | T8785C | L87P |
| MT-ATP6 | C8793G | H90D |
| MT-ATP6 | C8809T | T95I |

| | | |
|---|---|---|
| MT-ATP6 | C8814A | Q97K |
| MT-ATP6 | C8834A | *Syn* |
| MT-ATP6 | C8844A | P107T |
| MT-ATP6 | C8846A | *Syn* |
| MT-ATP6 | C8881A | S119Y |
| MT-ATP6 | C8881G | S119C |
| MT-ATP6 | G8885T | K120N |
| MT-ATP6 | A8889G | K122E |
| MT-ATP6 | C8897A | *Syn* |
| MT-ATP6 | C8898A | L125M |
| MT-ATP6 | C8898G | L125V |
| MT-ATP6 | G8901A | A126T |
| MT-ATP6 | G8901T | A126S |
| MT-ATP6 | A8905C | H127P |
| MT-ATP6 | A8912C | L129F |
| MT-ATP6 | C8914A | P130Q |
| MT-ATP6 | C8914T | P130L |
| MT-ATP6 | A8918G | *Syn* |
| MT-ATP6 | G8919A | G132S |
| MT-ATP6 | G8919C | G132R |
| MT-ATP6 | G8919T | G132C |
| MT-ATP6 | C8921A | *Syn* |
| MT-ATP6 | C8921T | *Syn* |
| MT-ATP6 | C8926A | P134H |
| MT-ATP6 | C8926T | P134L |
| MT-ATP6 | C8934A | L137I |
| MT-ATP6 | C8934T | L137F |
| MT-ATP6 | G8949T | V142F |
| MT-ATP6 | T8951C | *Syn* |
| MT-ATP6 | T8965A | I147N |
| MT-ATP6 | C8966A | I147M |
| MT-ATP6 | C8969G | S148Ter |
| MT-ATP6 | C8970A | L149M |
| MT-ATP6 | C8973A | L150I |
| MT-ATP6 | T8974C | L150P |
| MT-ATP6 | A8976G | I151V |
| MT-ATP6 | T8977C | I151T |
| MT-ATP6 | C8983A | P153Q |
| MT-ATP6 | T8986A | M154K |
| MT-ATP6 | G8988A | A155T |
| MT-ATP6 | G8988T | A155S |
| MT-ATP6 | C8990A | *Syn* |
| MT-ATP6 | G9001A | R159H |
| MT-ATP6 | G9001T | R159L |
| MT-ATP6 | C9002G | *Syn* |
| MT-ATP6 | C9003A | L160M |
| MT-ATP6 | T9020A | *Syn* |
| MT-ATP6 | C9027A | H168N |
| MT-ATP6 | C9027T | H168Y |
| MT-ATP6 | A9040C | H172P |
| MT-ATP6 | G9048C | G175R |
| MT-ATP6 | C9056A | *Syn* |
| MT-ATP6 | A9079C | N185T |
| MT-ATP6 | C9084A | P187T |
| MT-ATP6 | C9084T | P187S |

| | | |
|---|---|---|
| MT-ATP6 | C9084G | P187A |
| MT-ATP6 | T9087C | S188P |
| MT-ATP6 | C9111G | L196V |
| MT-ATP6 | C9117A | L198M |
| MT-ATP6 | C9117T | *Syn* |
| MT-ATP6 | T9127C | I201T |
| MT-ATP6 | C9137A | I204M |
| MT-ATP6 | G9144A | A207T |
| MT-ATP6 | G9144C | A207P |
| MT-ATP6 | G9144T | A207S |
| MT-ATP6 | C9146G | *Syn* |
| MT-ATP6 | T9147A | L208M |
| MT-ATP6 | T9151C | I209T |
| MT-ATP6 | C9158A | *Syn* |
| MT-ATP6 | C9158T | *Syn* |
| MT-ATP6 | G9177A | V218M |
| MT-ATP6 | C9183A | L220I |
| MT-ATP6 | A9187C | Y221S |
| MT-ATP6 | C9189A | L222M |
| MT-ATP6 | G9191A | *Syn* |
| MT-ATP6 | G9191T | *Syn* |
| MT-ATP6 | C9192A | H223N |
| MT-ATP6 | C9192T | H223Y |
| MT-ATP6 | G9195T | D224Y |
| MT-CO3 | A9213C | H3P |
| MT-CO3 | C9214A | H3Q |
| MT-CO3 | T9223A | H6Q |
| MT-CO3 | G9224A | A7T |
| MT-CO3 | G9224C | A7P |
| MT-CO3 | G9224T | A7S |
| MT-CO3 | C9226A | *Syn* |
| MT-CO3 | C9226T | *Syn* |
| MT-CO3 | T9229A | Y8Ter |
| MT-CO3 | A9235T | M10I |
| MT-CO3 | C9243A | P13H |
| MT-CO3 | C9244A | *Syn* |
| MT-CO3 | C9244T | *Syn* |
| MT-CO3 | C9244G | *Syn* |
| MT-CO3 | C9247A | S14Ter |
| MT-CO3 | C9248A | P15T |
| MT-CO3 | C9248T | P15S |
| MT-CO3 | T9251A | W16Ter |
| MT-CO3 | C9257A | L18M |
| MT-CO3 | C9261A | T19K |
| MT-CO3 | G9263A | G20Ter |
| MT-CO3 | G9263C | G20R |
| MT-CO3 | G9263T | G20W |
| MT-CO3 | G9265A | *Syn* |
| MT-CO3 | C9268A | *Syn* |
| MT-CO3 | C9269A | L22I |
| MT-CO3 | T9270A | L22H |
| MT-CO3 | T9270C | L22P |
| MT-CO3 | C9271G | *Syn* |
| MT-CO3 | T9282C | L26P |
| MT-CO3 | C9289T | *Syn* |

| | | |
|---|---|---|
| MT-CO3 | T9290A | S29T |
| MT-CO3 | C9291A | S29Y |
| MT-CO3 | G9294T | G30V |
| MT-CO3 | C9295A | *Syn* |
| MT-CO3 | C9300A | A32D |
| MT-CO3 | T9305C | W34R |
| MT-CO3 | C9316A | F37L |
| MT-CO3 | C9316T | *Syn* |
| MT-CO3 | C9317A | H38N |
| MT-CO3 | C9321A | S39Y |
| MT-CO3 | C9327T | T41M |
| MT-CO3 | C9332A | L43I |
| MT-CO3 | C9332T | L43F |
| MT-CO3 | C9338A | L45M |
| MT-CO3 | T9345A | L47Q |
| MT-CO3 | C9359T | *Syn* |
| MT-CO3 | C9370T | *Syn* |
| MT-CO3 | G9375C | W57S |
| MT-CO3 | G9375T | W57L |
| MT-CO3 | A9376G | *Syn* |
| MT-CO3 | T9377A | W58Ter |
| MT-CO3 | C9380A | R59S |
| MT-CO3 | G9386T | V61L |
| MT-CO3 | A9389C | T62P |
| MT-CO3 | C9402A | T66K |
| MT-CO3 | C9407A | Q68K |
| MT-CO3 | C9412A | *Syn* |
| MT-CO3 | C9420T | T72M |
| MT-CO3 | C9423A | P73Q |
| MT-CO3 | C9423G | P73R |
| MT-CO3 | C9425A | P74T |
| MT-CO3 | C9425T | P74S |
| MT-CO3 | T9427A | *Syn* |
| MT-CO3 | T9429C | V75A |
| MT-CO3 | C9430A | *Syn* |
| MT-CO3 | G9451A | *Syn* |
| MT-CO3 | T9453A | M83K |
| MT-CO3 | C9484A | F93L |
| MT-CO3 | T9485C | F94L |
| MT-CO3 | C9487A | F94L |
| MT-CO3 | C9487T | *Syn* |
| MT-CO3 | G9492T | G96V |
| MT-CO3 | G9503T | A100S |
| MT-CO3 | C9505A | *Syn* |
| MT-CO3 | C9511A | Y102Ter |
| MT-CO3 | C9517A | *Syn* |
| MT-CO3 | C9517G | *Syn* |
| MT-CO3 | C9520A | S105Ter |
| MT-CO3 | C9521T | *Syn* |
| MT-CO3 | A9530C | T109P |
| MT-CO3 | C9533A | P110T |
| MT-CO3 | A9541G | *Syn* |
| MT-CO3 | G9542T | G113W |
| MT-CO3 | G9545A | G114Ter |
| MT-CO3 | G9545T | G114W |

| | | |
|---|---|---|
| MT-CO3 | G9546A | G114E |
| MT-CO3 | G9552T | W116L |
| MT-CO3 | C9555A | P117H |
| MT-CO3 | A9559G | *Syn* |
| MT-CO3 | G9563T | G120C |
| MT-CO3 | C9582A | P126H |
| MT-CO3 | C9584A | L127M |
| MT-CO3 | G9587C | E128Q |
| MT-CO3 | C9592A | *Syn* |
| MT-CO3 | C9592T | *Syn* |
| MT-CO3 | C9596T | L131F |
| MT-CO3 | C9598A | *Syn* |
| MT-CO3 | C9598T | *Syn* |
| MT-CO3 | C9609T | S135F |
| MT-CO3 | C9610A | *Syn* |
| MT-CO3 | C9617T | L138F |
| MT-CO3 | C9619A | *Syn* |
| MT-CO3 | C9621A | A139E |
| MT-CO3 | C9621T | A139V |
| MT-CO3 | T9646A | *Syn* |
| MT-CO3 | C9650A | H149N |
| MT-CO3 | C9650T | H149Y |
| MT-CO3 | T9657A | L151Q |
| MT-CO3 | G9662T | E153Ter |
| MT-CO3 | C9670T | *Syn* |
| MT-CO3 | C9686G | Q161E |
| MT-CO3 | G9694T | *Syn* |
| MT-CO3 | C9695A | L164I |
| MT-CO3 | G9713T | G170C |
| MT-CO3 | C9718T | *Syn* |
| MT-CO3 | C9731A | L176M |
| MT-CO3 | T9757A | *Syn* |
| MT-CO3 | C9760A | *Syn* |
| MT-CO3 | T9761G | F186V |
| MT-CO3 | T9762A | F186Y |
| MT-CO3 | C9772A | *Syn* |
| MT-CO3 | C9775T | *Syn* |
| MT-CO3 | G9776C | G191R |
| MT-CO3 | G9776T | G191C |
| MT-CO3 | G9785A | G194S |
| MT-CO3 | G9785T | G194C |
| MT-CO3 | C9787A | *Syn* |
| MT-CO3 | C9787T | *Syn* |
| MT-CO3 | T9788A | S195T |
| MT-CO3 | C9804A | A200D |
| MT-CO3 | C9805A | *Syn* |
| MT-CO3 | G9809T | G202C |
| MT-CO3 | T9812C | F203L |
| MT-CO3 | C9815A | H204N |
| MT-CO3 | C9815T | H204Y |
| MT-CO3 | A9816C | H204P |
| MT-CO3 | G9818A | G205Ter |
| MT-CO3 | G9818C | G205R |
| MT-CO3 | G9818T | G205W |
| MT-CO3 | G9819T | G205V |

| | | |
|---|---|---|
| MT-CO3 | A9820C | *Syn* |
| MT-CO3 | T9822A | L206H |
| MT-CO3 | T9823A | *Syn* |
| MT-CO3 | C9824A | H207N |
| MT-CO3 | C9824T | H207Y |
| MT-CO3 | C9824G | H207D |
| MT-CO3 | A9830T | I209F |
| MT-CO3 | C9843T | T213I |
| MT-CO3 | C9847T | *Syn* |
| MT-CO3 | C9850A | *Syn* |
| MT-CO3 | A9851C | T216P |
| MT-CO3 | C9852A | T216N |
| MT-CO3 | C9856A | I217M |
| MT-CO3 | C9856T | *Syn* |
| MT-CO3 | T9861A | F219Y |
| MT-CO3 | T9861C | F219S |
| MT-CO3 | C9866A | R221S |
| MT-CO3 | C9866G | R221G |
| MT-CO3 | C9869T | Q222Ter |
| MT-CO3 | C9891A | S229Y |
| MT-CO3 | C9899A | H232N |
| MT-CO3 | C9901T | *Syn* |
| MT-CO3 | C9915A | A237D |
| MT-CO3 | C9915T | A237V |
| MT-CO3 | C9916A | *Syn* |
| MT-CO3 | C9922A | *Syn* |
| MT-CO3 | A9927G | Y241C |
| MT-CO3 | G9938T | V245L |
| MT-CO3 | G9944T | V247L |
| MT-CO3 | C9953A | L250M |
| MT-CO3 | T9958G | F251L |
| MT-CO3 | G9961T | *Syn* |
| MT-CO3 | T9964C | *Syn* |
| MT-CO3 | C9967A | *Syn* |
| MT-CO3 | C9973A | I256M |
| MT-CO3 | G9981T | W259L |
| MT-CO3 | T9986G | S261A |
| MT-ND3 | A10078T | L7F |
| MT-ND3 | C10084A | I9M |
| MT-ND3 | C10093A | *Syn* |
| MT-ND3 | C10094T | *Syn* |
| MT-ND3 | C10098A | A14D |
| MT-ND3 | C10103A | L16M |
| MT-ND3 | T10104C | L16P |
| MT-ND3 | A10105T | *Syn* |
| MT-ND3 | T10107A | L17Q |
| MT-ND3 | A10112T | I19F |
| MT-ND3 | A10118G | T21A |
| MT-ND3 | C10130A | P25T |
| MT-ND3 | C10130T | P25S |
| MT-ND3 | C10131A | P25Q |
| MT-ND3 | A10135C | Q26H |
| MT-ND3 | C10144A | *Syn* |
| MT-ND3 | T10157C | S34P |
| MT-ND3 | C10158A | S34Y |

| | | |
|---|---|---|
| MT-ND3 | C10158G | S34C |
| MT-ND3 | C10159A | *Syn* |
| MT-ND3 | C10159G | *Syn* |
| MT-ND3 | C10164A | P36H |
| MT-ND3 | C10164T | P36L |
| MT-ND3 | A10167G | Y37C |
| MT-ND3 | C10168T | *Syn* |
| MT-ND3 | G10171A | *Syn* |
| MT-ND3 | G10171T | E38D |
| MT-ND3 | C10174T | *Syn* |
| MT-ND3 | T10190C | S45P |
| MT-ND3 | C10191T | S45F |
| MT-ND3 | C10192A | *Syn* |
| MT-ND3 | C10195A | *Syn* |
| MT-ND3 | C10195T | *Syn* |
| MT-ND3 | C10195G | *Syn* |
| MT-ND3 | C10201T | *Syn* |
| MT-ND3 | G10229A | V58M |
| MT-ND3 | C10233T | A59V |
| MT-ND3 | C10240A | *Syn* |
| MT-ND3 | G10253A | D66N |
| MT-ND3 | G10253T | D66Y |
| MT-ND3 | C10361A | L102M |
| MT-ND3 | C10365A | A103D |
| MT-ND3 | C10366A | *Syn* |
| MT-ND3 | A10396G | *Syn* |
| MT-ND3 | A10397G | T114A |
| MT-ND4L | C10472A | P2T |
| MT-ND4L | C10473A | P2H |
| MT-ND4L | C10475A | L3I |
| MT-ND4L | C10475T | L3F |
| MT-ND4L | C10477A | *Syn* |
| MT-ND4L | G10499T | A11S |
| MT-ND4L | C10506A | T13N |
| MT-ND4L | C10506T | T13I |
| MT-ND4L | T10511C | S15P |
| MT-ND4L | C10512A | S15Ter |
| MT-ND4L | C10539A | S24Ter |
| MT-ND4L | C10539G | S24W |
| MT-ND4L | A10542C | H25P |
| MT-ND4L | C10543T | *Syn* |
| MT-ND4L | C10552A | *Syn* |
| MT-ND4L | T10553C | S29P |
| MT-ND4L | C10565A | L33M |
| MT-ND4L | C10584A | S39Ter |
| MT-ND4L | C10584T | S39L |
| MT-ND4L | C10584G | S39W |
| MT-ND4L | C10591A | F41L |
| MT-ND4L | G10598A | A44T |
| MT-ND4L | C10599A | A44D |
| MT-ND4L | C10599T | A44V |
| MT-ND4L | C10602T | T45I |
| MT-ND4L | T10603A | *Syn* |
| MT-ND4L | C10604A | L46I |
| MT-ND4L | T10605A | L46H |

| | | |
|---|---|---|
| MT-ND4L | T10614C | L49P |
| MT-ND4L | A10619C | T51P |
| MT-ND4L | T10625A | S53T |
| MT-ND4L | C10630A | *Syn* |
| MT-ND4L | C10635A | A56D |
| MT-ND4L | C10635T | A56V |
| MT-ND4L | T10639C | *Syn* |
| MT-ND4L | T10651A | I61M |
| MT-ND4L | G10652T | A62S |
| MT-ND4L | C10653A | A62D |
| MT-ND4L | C10653T | A62V |
| MT-ND4L | A10655T | M63L |
| MT-ND4L | A10657G | *Syn* |
| MT-ND4L | C10668T | A67V |
| MT-ND4L | C10669A | *Syn* |
| MT-ND4L | C10672A | *Syn* |
| MT-ND4L | G10688A | G74S |
| MT-ND4L | G10689A | G74D |
| MT-ND4L | C10696A | *Syn* |
| MT-ND4L | C10705A | *Syn* |
| MT-ND4L | C10714A | *Syn* |
| MT-ND4L | A10716G | N83S |
| MT-ND4L | C10717T | *Syn* |
| MT-ND4L | C10719A | T84K |
| MT-ND4L | G10724A | G86S |
| MT-ND4L | G10724T | G86C |
| MT-ND4L | C10726A | *Syn* |
| MT-ND4L | C10726T | *Syn* |
| MT-ND4L | G10736A | V90M |
| MT-ND4L | G10736T | V90L |
| MT-ND4L | A10740T | H91L |
| MT-ND4L | C10744A | N92K |
| MT-ND4L | C10751A | L95M |
| MT-ND4L | C10751G | L95V |
| MT-ND4L | C10756A | *Syn* |
| MT-ND4L | C10757A | Q97K |
| MT-ND4L | T10760A | C98S |
| MT-ND4 | T10760A | M1K |
| MT-ND4L | G10761T | C98F |
| MT-ND4 | G10761T | M1I |
| MT-ND4 | C10768A | L4M |
| MT-ND4 | C10773A | I5M |
| MT-ND4 | C10796A | P13Q |
| MT-ND4 | C10796T | P13L |
| MT-ND4 | C10796G | P13R |
| MT-ND4 | G10800A | *Syn* |
| MT-ND4 | G10800C | *Syn* |
| MT-ND4 | G10800T | *Syn* |
| MT-ND4 | G10805T | W16L |
| MT-ND4 | C10819A | H21N |
| MT-ND4 | C10819T | H21Y |
| MT-ND4 | T10826C | I23T |
| MT-ND4 | C10836T | *Syn* |
| MT-ND4 | A10839G | *Syn* |
| MT-ND4 | A10840C | T28P |

| | | |
|---|---|---|
| MT-ND4 | C10842A | *Syn* |
| MT-ND4 | C10845A | *Syn* |
| MT-ND4 | C10845G | *Syn* |
| MT-ND4 | C10848A | H30Q |
| MT-ND4 | C10852T | *Syn* |
| MT-ND4 | A10855T | I33F |
| MT-ND4 | T10859C | I34T |
| MT-ND4 | G10862A | S35N |
| MT-ND4 | G10862T | S35I |
| MT-ND4 | C10863A | S35Ter |
| MT-ND4 | C10866A | I36M |
| MT-ND4 | A10867C | I37L |
| MT-ND4 | A10867G | I37V |
| MT-ND4 | C10876A | L40M |
| MT-ND4 | T10892A | I45N |
| MT-ND4 | C10903A | L49M |
| MT-ND4 | G10913A | C52Y |
| MT-ND4 | C10922A | T55N |
| MT-ND4 | C10931A | S58Y |
| MT-ND4 | C10931T | S58F |
| MT-ND4 | G10933C | D59H |
| MT-ND4 | G10933T | D59Y |
| MT-ND4 | T10940C | L61P |
| MT-ND4 | C10946A | T63N |
| MT-ND4 | C10948A | P64T |
| MT-ND4 | C10950A | *Syn* |
| MT-ND4 | C10950T | *Syn* |
| MT-ND4 | C10951A | L65I |
| MT-ND4 | C10953A | *Syn* |
| MT-ND4 | T10955C | L66P |
| MT-ND4 | C10967T | T70I |
| MT-ND4 | C10968A | *Syn* |
| MT-ND4 | G10970T | W71L |
| MT-ND4 | T10973C | L72P |
| MT-ND4 | C10974A | *Syn* |
| MT-ND4 | A10977C | *Syn* |
| MT-ND4 | A10977G | *Syn* |
| MT-ND4 | C10979A | P74H |
| MT-ND4 | G10992T | M78I |
| MT-ND4 | G10993A | A79T |
| MT-ND4 | G10993T | A79S |
| MT-ND4 | C10994T | A79V |
| MT-ND4 | A11014G | S86G |
| MT-ND4 | C11021A | P88Q |
| MT-ND4 | C11023T | *Syn* |
| MT-ND4 | C11027A | S90Ter |
| MT-ND4 | C11040A | *Syn* |
| MT-ND4 | C11046A | *Syn* |
| MT-ND4 | C11048A | S97Y |
| MT-ND4 | C11060T | S101F |
| MT-ND4 | C11061A | *Syn* |
| MT-ND4 | T11071C | S105P |
| MT-ND4 | C11073A | *Syn* |
| MT-ND4 | C11073T | *Syn* |
| MT-ND4 | C11073G | *Syn* |

| | | |
|---|---|---|
| MT-ND4 | A11085T | *Syn* |
| MT-ND4 | C11093A | A112D |
| MT-ND4 | A11109T | M117I |
| MT-ND4 | A11114T | Y119F |
| MT-ND4 | C11118T | *Syn* |
| MT-ND4 | C11132T | T125M |
| MT-ND4 | T11135A | L126H |
| MT-ND4 | C11139A | I127M |
| MT-ND4 | C11140A | P128T |
| MT-ND4 | C11140G | P128A |
| MT-ND4 | G11148T | L130F |
| MT-ND4 | C11154A | I132M |
| MT-ND4 | C11157A | I133M |
| MT-ND4 | C11160A | *Syn* |
| MT-ND4 | C11160G | *Syn* |
| MT-ND4 | C11161A | R135Ter |
| MT-ND4 | G11167A | G137S |
| MT-ND4 | G11167T | G137C |
| MT-ND4 | C11173A | Q139K |
| MT-ND4 | G11179A | E141K |
| MT-ND4 | C11182A | R142S |
| MT-ND4 | C11182T | R142C |
| MT-ND4 | C11185A | L143M |
| MT-ND4 | C11196A | *Syn* |
| MT-ND4 | C11202T | *Syn* |
| MT-ND4 | C11211A | F151L |
| MT-ND4 | C11216A | T153N |
| MT-ND4 | C11230G | L158V |
| MT-ND4 | C11234A | P159H |
| MT-ND4 | C11235A | *Syn* |
| MT-ND4 | C11236A | L160M |
| MT-ND4 | C11244G | I162M |
| MT-ND4 | C11246A | A163E |
| MT-ND4 | C11262A | H168Q |
| MT-ND4 | A11264G | N169S |
| MT-ND4 | C11268A | *Syn* |
| MT-ND4 | C11269A | L171M |
| MT-ND4 | C11269T | *Syn* |
| MT-ND4 | T11270C | L171P |
| MT-ND4 | A11271T | *Syn* |
| MT-ND4 | C11278T | *Syn* |
| MT-ND4 | C11283A | N175K |
| MT-ND4 | T11294A | L179H |
| MT-ND4 | C11295A | *Syn* |
| MT-ND4 | C11347A | L197M |
| MT-ND4 | C11347T | *Syn* |
| MT-ND4 | T11348C | L197P |
| MT-ND4 | G11350T | A198S |
| MT-ND4 | C11351A | A198D |
| MT-ND4 | C11351T | A198V |
| MT-ND4 | C11357A | T200K |
| MT-ND4 | A11358T | *Syn* |
| MT-ND4 | A11359T | M201L |
| MT-ND4 | G11371T | V205L |
| MT-ND4 | C11380A | P208T |

| | | |
|---|---|---|
| MT-ND4 | G11389T | G211W |
| MT-ND4 | G11390T | G211V |
| MT-ND4 | C11394A | *Syn* |
| MT-ND4 | G11402T | W215L |
| MT-ND4 | C11404A | L216I |
| MT-ND4 | C11407A | P217T |
| MT-ND4 | C11414T | A219V |
| MT-ND4 | C11428A | P224T |
| MT-ND4 | T11432C | I225T |
| MT-ND4 | G11434T | A226S |
| MT-ND4 | C11435A | A226D |
| MT-ND4 | A11445G | *Syn* |
| MT-ND4 | C11453A | A232D |
| MT-ND4 | C11454A | *Syn* |
| MT-ND4 | C11454T | *Syn* |
| MT-ND4 | C11461T | L235F |
| MT-ND4 | C11463A | *Syn* |
| MT-ND4 | T11464G | L236V |
| MT-ND4 | G11473A | G239S |
| MT-ND4 | C11475T | *Syn* |
| MT-ND4 | G11482C | G242R |
| MT-ND4 | G11482T | G242C |
| MT-ND4 | T11484G | *Syn* |
| MT-ND4 | C11491T | R245C |
| MT-ND4 | C11493A | *Syn* |
| MT-ND4 | C11493T | *Syn* |
| MT-ND4 | C11502A | *Syn* |
| MT-ND4 | C11513A | P252H |
| MT-ND4 | C11515A | L253M |
| MT-ND4 | C11524T | H256Y |
| MT-ND4 | T11539C | F261L |
| MT-ND4 | G11545T | V263L |
| MT-ND4 | C11548T | *Syn* |
| MT-ND4 | C11553A | *Syn* |
| MT-ND4 | G11561T | G268V |
| MT-ND4 | C11562T | *Syn* |
| MT-ND4 | T11570A | M271K |
| MT-ND4 | G11576T | S273I |
| MT-ND4 | C11579A | S274Y |
| MT-ND4 | T11584A | C276S |
| MT-ND4 | T11588A | L277Q |
| MT-ND4 | C11590G | R278G |
| MT-ND4 | C11593G | Q279E |
| MT-ND4 | G11610T | *Syn* |
| MT-ND4 | C11613A | *Syn* |
| MT-ND4 | C11618T | A287V |
| MT-ND4 | C11622A | Y288Ter |
| MT-ND4 | C11622G | Y288Ter |
| MT-ND4 | G11633A | S292N |
| MT-ND4 | G11633T | S292I |
| MT-ND4 | C11637A | H293Q |
| MT-ND4 | C11644A | L296I |
| MT-ND4 | C11644G | L296V |
| MT-ND4 | G11650A | V298M |
| MT-ND4 | C11658A | *Syn* |

| | | |
|---|---|---|
| MT-ND4 | C11658G | *Syn* |
| MT-ND4 | C11662A | L302I |
| MT-ND4 | C11664A | *Syn* |
| MT-ND4 | C11664T | *Syn* |
| MT-ND4 | C11668A | Q304K |
| MT-ND4 | C11668G | Q304E |
| MT-ND4 | C11675A | P306H |
| MT-ND4 | C11676A | *Syn* |
| MT-ND4 | C11691A | *Syn* |
| MT-ND4 | C11691G | *Syn* |
| MT-ND4 | C11713A | H319N |
| MT-ND4 | G11717T | G320V |
| MT-ND4 | C11726A | S323Y |
| MT-ND4 | C11726G | S323C |
| MT-ND4 | G11741A | C328Y |
| MT-ND4 | G11741T | C328F |
| MT-ND4 | G11746T | A330S |
| MT-ND4 | C11757A | N333K |
| MT-ND4 | T11758A | Y334N |
| MT-ND4 | C11768A | T337N |
| MT-ND4 | C11768T | T337I |
| MT-ND4 | C11772A | H338Q |
| MT-ND4 | G11774A | S339N |
| MT-ND4 | C11776A | R340S |
| MT-ND4 | G11777A | R340H |
| MT-ND4 | G11777T | R340L |
| MT-ND4 | C11778T | *Syn* |
| MT-ND4 | C11792A | S345Y |
| MT-ND4 | G11798C | G347A |
| MT-ND4 | G11798T | G347V |
| MT-ND4 | C11800A | L348I |
| MT-ND4 | C11809A | L351M |
| MT-ND4 | G11824A | A356T |
| MT-ND4 | G11824T | A356S |
| MT-ND4 | C11825T | A356V |
| MT-ND4 | A11832T | W358C |
| MT-ND4 | C11836T | L360F |
| MT-ND4 | C11843A | A362E |
| MT-ND4 | G11846A | S363N |
| MT-ND4 | G11846T | S363I |
| MT-ND4 | C11848A | L364I |
| MT-ND4 | C11857A | L367I |
| MT-ND4 | C11859A | *Syn* |
| MT-ND4 | G11860C | A368P |
| MT-ND4 | G11860T | A368S |
| MT-ND4 | C11862A | *Syn* |
| MT-ND4 | C11862T | *Syn* |
| MT-ND4 | C11862G | *Syn* |
| MT-ND4 | C11869A | P371T |
| MT-ND4 | C11869T | P371S |
| MT-ND4 | C11871A | *Syn* |
| MT-ND4 | C11871G | *Syn* |
| MT-ND4 | C11873T | T372I |
| MT-ND4 | C11880A | N374K |
| MT-ND4 | C11880T | *Syn* |

| | | |
|---|---|---|
| MT-ND4 | G11890T | E378Ter |
| MT-ND4 | C11893T | L379F |
| MT-ND4 | T11898C | *Syn* |
| MT-ND4 | G11899A | V381M |
| MT-ND4 | C11902A | L382M |
| MT-ND4 | C11910A | *Syn* |
| MT-ND4 | C11918G | S387C |
| MT-ND4 | C11919A | *Syn* |
| MT-ND4 | T11920A | W388Ter |
| MT-ND4 | A11927T | N390I |
| MT-ND4 | T11928C | *Syn* |
| MT-ND4 | C11931A | I391M |
| MT-ND4 | C11931G | I391M |
| MT-ND4 | A11932C | T392P |
| MT-ND4 | C11933A | T392N |
| MT-ND4 | C11941A | L395I |
| MT-ND4 | G11948T | G397V |
| MT-ND4 | C11955A | N399K |
| MT-ND4 | A11956T | M400L |
| MT-ND4 | C11964A | *Syn* |
| MT-ND4 | C11971A | L405M |
| MT-ND4 | C11976T | *Syn* |
| MT-ND4 | C11980A | L408I |
| MT-ND4 | C12001T | Q415Ter |
| MT-ND4 | G12005A | W416Ter |
| MT-ND4 | G12005C | W416S |
| MT-ND4 | G12005T | W416L |
| MT-ND4 | C12011A | S418Ter |
| MT-ND4 | C12011T | S418L |
| MT-ND4 | C12017A | T420N |
| MT-ND4 | C12017T | T420I |
| MT-ND4 | C12018A | *Syn* |
| MT-ND4 | C12018G | *Syn* |
| MT-ND4 | C12022A | H422N |
| MT-ND4 | C12022T | H422Y |
| MT-ND4 | A12025T | I423F |
| MT-ND4 | C12042A | *Syn* |
| MT-ND4 | C12052A | R432Ter |
| MT-ND4 | G12055T | E433Ter |
| MT-ND4 | C12060T | *Syn* |
| MT-ND4 | A12061C | T435P |
| MT-ND4 | C12062A | T435N |
| MT-ND4 | T12065C | L436P |
| MT-ND4 | C12072T | *Syn* |
| MT-ND4 | T12074C | M439T |
| MT-ND4 | C12079A | L441M |
| MT-ND4 | T12082A | S442T |
| MT-ND4 | C12083A | S442Y |
| MT-ND4 | C12083T | S442F |
| MT-ND4 | C12083G | S442C |
| MT-ND4 | C12094A | L446I |
| MT-ND4 | C12094T | L446F |
| MT-ND4 | C12110A | P451H |
| MT-ND4 | C12117A | I453M |
| MT-ND4 | C12122T | T455I |

| | | |
|---|---|---|
| MT-ND4 | T12128C | F457S |
| MT-ND4 | C12132A | *Syn* |
| MT-ND4 | T12136A | C460S |
| MT-ND5 | T12337A | M1K |
| MT-ND5 | C12340A | T2N |
| MT-ND5 | T12343C | M3T |
| MT-ND5 | C12345A | H4N |
| MT-ND5 | C12347A | H4Q |
| MT-ND5 | C12349A | T5N |
| MT-ND5 | A12351G | T6A |
| MT-ND5 | C12352A | T6N |
| MT-ND5 | C12352T | T6I |
| MT-ND5 | T12364C | L10P |
| MT-ND5 | A12365G | *Syn* |
| MT-ND5 | C12368A | *Syn* |
| MT-ND5 | C12373A | T13N |
| MT-ND5 | C12376T | S14F |
| MT-ND5 | C12377A | *Syn* |
| MT-ND5 | C12377T | *Syn* |
| MT-ND5 | C12377G | *Syn* |
| MT-ND5 | C12378A | L15M |
| MT-ND5 | C12378T | *Syn* |
| MT-ND5 | C12384A | P17T |
| MT-ND5 | C12384T | P17S |
| MT-ND5 | C12410T | *Syn* |
| MT-ND5 | C12411A | P26T |
| MT-ND5 | C12448A | S38Y |
| MT-ND5 | T12452A | I39M |
| MT-ND5 | C12455A | *Syn* |
| MT-ND5 | C12455T | *Syn* |
| MT-ND5 | C12457T | A41V |
| MT-ND5 | T12465C | F44L |
| MT-ND5 | C12477A | L48I |
| MT-ND5 | T12481C | F49S |
| MT-ND5 | C12482A | F49L |
| MT-ND5 | C12484A | P50H |
| MT-ND5 | A12486C | T51P |
| MT-ND5 | C12497A | F54L |
| MT-ND5 | T12501A | C56S |
| MT-ND5 | G12502A | C56Y |
| MT-ND5 | G12502T | C56F |
| MT-ND5 | C12503A | C56W |
| MT-ND5 | G12507A | D58N |
| MT-ND5 | G12507C | D58H |
| MT-ND5 | G12507T | D58Y |
| MT-ND5 | C12510A | Q59K |
| MT-ND5 | C12510G | Q59E |
| MT-ND5 | A12533G | *Syn* |
| MT-ND5 | C12542A | *Syn* |
| MT-ND5 | C12548A | *Syn* |
| MT-ND5 | C12549A | Q72K |
| MT-ND5 | C12565T | S77F |
| MT-ND5 | C12565G | S77C |
| MT-ND5 | C12567A | L78M |
| MT-ND5 | T12568C | L78P |

| | | |
|---|---|---|
| MT-ND5 | C12572A | S79Ter |
| MT-ND5 | G12582A | D83N |
| MT-ND5 | G12582C | D83H |
| MT-ND5 | G12582T | D83Y |
| MT-ND5 | C12584T | *Syn* |
| MT-ND5 | T12589C | F85S |
| MT-ND5 | C12592A | S86Y |
| MT-ND5 | A12603G | I90V |
| MT-ND5 | C12606A | P91T |
| MT-ND5 | C12632A | *Syn* |
| MT-ND5 | C12644A | F103L |
| MT-ND5 | T12649C | L105P |
| MT-ND5 | G12650A | *Syn* |
| MT-ND5 | G12652A | W106Ter |
| MT-ND5 | G12652T | W106L |
| MT-ND5 | G12666T | D111Y |
| MT-ND5 | A12667C | D111A |
| MT-ND5 | C12670A | P112Q |
| MT-ND5 | C12681A | Q116K |
| MT-ND5 | T12695C | *Syn* |
| MT-ND5 | C12699T | L122F |
| MT-ND5 | A12710T | *Syn* |
| MT-ND5 | C12715T | T127I |
| MT-ND5 | C12716A | *Syn* |
| MT-ND5 | T12718A | M128K |
| MT-ND5 | G12729A | V132I |
| MT-ND5 | C12734A | *Syn* |
| MT-ND5 | C12734T | *Syn* |
| MT-ND5 | C12744A | L137M |
| MT-ND5 | T12748C | F138S |
| MT-ND5 | C12749A | F138L |
| MT-ND5 | G12755T | *Syn* |
| MT-ND5 | C12758A | F141L |
| MT-ND5 | C12761A | I142M |
| MT-ND5 | G12770T | E145D |
| MT-ND5 | C12773A | *Syn* |
| MT-ND5 | C12773T | *Syn* |
| MT-ND5 | C12773G | *Syn* |
| MT-ND5 | G12777T | G148W |
| MT-ND5 | C12787A | S151Y |
| MT-ND5 | C12788A | *Syn* |
| MT-ND5 | T12790C | F152S |
| MT-ND5 | T12799A | I155N |
| MT-ND5 | T12799C | I155T |
| MT-ND5 | C12800A | I155M |
| MT-ND5 | C12800G | I155M |
| MT-ND5 | G12808T | W158L |
| MT-ND5 | C12815A | *Syn* |
| MT-ND5 | C12815T | *Syn* |
| MT-ND5 | A12823T | D163V |
| MT-ND5 | C12827A | *Syn* |
| MT-ND5 | A12840T | I169F |
| MT-ND5 | C12843A | Q170K |
| MT-ND5 | G12846A | A171T |
| MT-ND5 | A12848T | *Syn* |

| | | |
|---|---|---|
| MT-ND5 | C12851A | I172M |
| MT-ND5 | C12851T | *Syn* |
| MT-ND5 | C12860A | N175K |
| MT-ND5 | G12867A | G178S |
| MT-ND5 | G12867T | G178C |
| MT-ND5 | G12870T | D179Y |
| MT-ND5 | C12875A | I180M |
| MT-ND5 | T12878G | *Syn* |
| MT-ND5 | C12887A | *Syn* |
| MT-ND5 | C12887G | *Syn* |
| MT-ND5 | C12890A | *Syn* |
| MT-ND5 | C12890T | *Syn* |
| MT-ND5 | A12893G | *Syn* |
| MT-ND5 | G12894T | A187S |
| MT-ND5 | T12901A | F189Y |
| MT-ND5 | C12905A | I190M |
| MT-ND5 | C12905T | *Syn* |
| MT-ND5 | C12911A | H192Q |
| MT-ND5 | C12913A | S193Y |
| MT-ND5 | C12919A | S195Ter |
| MT-ND5 | A12920G | *Syn* |
| MT-ND5 | G12922C | W196S |
| MT-ND5 | C12926T | *Syn* |
| MT-ND5 | C12958A | P208Q |
| MT-ND5 | C12958T | P208L |
| MT-ND5 | C12963A | L210I |
| MT-ND5 | G12978T | G215C |
| MT-ND5 | C12980A | *Syn* |
| MT-ND5 | C12980T | *Syn* |
| MT-ND5 | C12981T | L216F |
| MT-ND5 | C12991T | A219V |
| MT-ND5 | G12993A | A220T |
| MT-ND5 | G12999A | G222S |
| MT-ND5 | G13000A | G222D |
| MT-ND5 | C13001A | *Syn* |
| MT-ND5 | C13001T | *Syn* |
| MT-ND5 | T13005C | S224P |
| MT-ND5 | C13009A | A225D |
| MT-ND5 | C13009T | A225V |
| MT-ND5 | G13017T | G228C |
| MT-ND5 | C13025A | H230Q |
| MT-ND5 | C13025T | *Syn* |
| MT-ND5 | C13028A | *Syn* |
| MT-ND5 | C13032A | L233I |
| MT-ND5 | C13034A | *Syn* |
| MT-ND5 | C13034T | *Syn* |
| MT-ND5 | C13035A | P234T |
| MT-ND5 | C13035T | P234S |
| MT-ND5 | C13037A | *Syn* |
| MT-ND5 | C13042A | A236D |
| MT-ND5 | G13050A | G239S |
| MT-ND5 | G13050T | G239C |
| MT-ND5 | G13051T | G239V |
| MT-ND5 | C13055A | *Syn* |
| MT-ND5 | C13064A | *Syn* |

| | | |
|---|---|---|
| MT-ND5 | C13064T | *Syn* |
| MT-ND5 | T13065C | S244P |
| MT-ND5 | C13070A | *Syn* |
| MT-ND5 | T13075C | L247P |
| MT-ND5 | C13077A | H248N |
| MT-ND5 | C13077T | H248Y |
| MT-ND5 | A13078G | H248R |
| MT-ND5 | C13079T | *Syn* |
| MT-ND5 | A13083T | S250C |
| MT-ND5 | G13095A | V254M |
| MT-ND5 | G13095T | V254L |
| MT-ND5 | C13099A | A255E |
| MT-ND5 | C13099T | A255V |
| MT-ND5 | G13102T | G256V |
| MT-ND5 | C13106A | I257M |
| MT-ND5 | C13109A | F258L |
| MT-ND5 | C13109T | *Syn* |
| MT-ND5 | C13113T | L260F |
| MT-ND5 | C13119A | R262S |
| MT-ND5 | T13122C | F263L |
| MT-ND5 | C13127G | H264Q |
| MT-ND5 | C13128A | P265T |
| MT-ND5 | C13129A | P265H |
| MT-ND5 | C13146A | P271T |
| MT-ND5 | C13146T | P271S |
| MT-ND5 | T13153A | I273N |
| MT-ND5 | T13160C | *Syn* |
| MT-ND5 | C13161T | *Syn* |
| MT-ND5 | C13165A | T277K |
| MT-ND5 | C13167A | L278M |
| MT-ND5 | G13171T | C279F |
| MT-ND5 | C13172A | C279W |
| MT-ND5 | G13177A | G281D |
| MT-ND5 | C13178A | *Syn* |
| MT-ND5 | T13183A | I283N |
| MT-ND5 | C13186A | T284N |
| MT-ND5 | C13186G | T284S |
| MT-ND5 | C13189T | T285I |
| MT-ND5 | C13191A | L286M |
| MT-ND5 | G13193A | *Syn* |
| MT-ND5 | G13200A | A289T |
| MT-ND5 | C13201G | A289G |
| MT-ND5 | G13203A | V290I |
| MT-ND5 | C13205A | *Syn* |
| MT-ND5 | C13205T | *Syn* |
| MT-ND5 | T13206A | C291S |
| MT-ND5 | C13211A | *Syn* |
| MT-ND5 | C13212A | L293I |
| MT-ND5 | C13212T | L293F |
| MT-ND5 | C13216A | T294K |
| MT-ND5 | C13216T | T294M |
| MT-ND5 | T13223A | N296K |
| MT-ND5 | T13223C | *Syn* |
| MT-ND5 | T13228A | I298N |
| MT-ND5 | T13228C | I298T |

| | | |
|---|---|---|
| MT-ND5 | G13239C | V302L |
| MT-ND5 | G13239T | V302L |
| MT-ND5 | C13243A | A303D |
| MT-ND5 | C13244A | *Syn* |
| MT-ND5 | C13244T | *Syn* |
| MT-ND5 | C13244G | *Syn* |
| MT-ND5 | A13251C | T306P |
| MT-ND5 | C13255A | S307Ter |
| MT-ND5 | G13258A | S308N |
| MT-ND5 | G13267A | G311E |
| MT-ND5 | G13267C | G311A |
| MT-ND5 | G13278T | V315F |
| MT-ND5 | C13286A | I317M |
| MT-ND5 | C13286T | *Syn* |
| MT-ND5 | C13286G | I317M |
| MT-ND5 | T13291A | I319N |
| MT-ND5 | C13295G | N320K |
| MT-ND5 | A13298G | *Syn* |
| MT-ND5 | T13311C | F326L |
| MT-ND5 | C13327T | T331I |
| MT-ND5 | C13328A | *Syn* |
| MT-ND5 | C13329A | H332N |
| MT-ND5 | C13329T | H332Y |
| MT-ND5 | A13330T | H332L |
| MT-ND5 | G13332C | A333P |
| MT-ND5 | G13332T | A333S |
| MT-ND5 | C13334A | *Syn* |
| MT-ND5 | C13334T | *Syn* |
| MT-ND5 | T13336C | F334S |
| MT-ND5 | C13340A | F335L |
| MT-ND5 | C13340T | *Syn* |
| MT-ND5 | C13340G | F335L |
| MT-ND5 | A13343G | *Syn* |
| MT-ND5 | G13344T | A337S |
| MT-ND5 | C13345A | A337D |
| MT-ND5 | C13350A | L339M |
| MT-ND5 | C13350T | *Syn* |
| MT-ND5 | C13363A | S343Y |
| MT-ND5 | C13379A | H348Q |
| MT-ND5 | C13388A | N351K |
| MT-ND5 | C13395T | Q354Ter |
| MT-ND5 | G13413A | G360Ter |
| MT-ND5 | G13413C | G360R |
| MT-ND5 | T13453C | L373P |
| MT-ND5 | C13467A | L378M |
| MT-ND5 | G13470T | A379S |
| MT-ND5 | C13617A | L428I |
| MT-ND5 | C13622A | *Syn* |
| MT-ND5 | G13633T | G433V |
| MT-ND5 | A13636G | Q434R |
| MT-ND5 | C13639A | P435H |
| MT-ND5 | C13639T | P435L |
| MT-ND5 | C13646A | F437L |
| MT-ND5 | C13652A | *Syn* |
| MT-ND5 | C13657A | T441N |

| | | |
|---|---|---|
| MT-ND5 | C13657T | T441I |
| MT-ND5 | A13660G | N442S |
| MT-ND5 | G13668T | E445Ter |
| MT-ND5 | C13677T | P448S |
| MT-ND5 | C13679A | *Syn* |
| MT-ND5 | G13706T | *Syn* |
| MT-ND5 | C13712A | *Syn* |
| MT-ND5 | C13719A | L462M |
| MT-ND5 | G13725T | A464S |
| MT-ND5 | G13729T | G465V |
| MT-ND5 | C13736A | *Syn* |
| MT-ND5 | A13737T | I468F |
| MT-ND5 | C13745T | *Syn* |
| MT-ND5 | C13754A | *Syn* |
| MT-ND5 | C13754T | *Syn* |
| MT-ND5 | C13762A | S476Y |
| MT-ND5 | C13763A | *Syn* |
| MT-ND5 | C13769T | *Syn* |
| MT-ND5 | C13770A | Q479K |
| MT-ND5 | C13781A | I482M |
| MT-ND5 | T13786C | L484P |
| MT-ND5 | C13790T | *Syn* |
| MT-ND5 | C13797A | L488I |
| MT-ND5 | C13797T | L488F |
| MT-ND5 | C13799A | *Syn* |
| MT-ND5 | C13799G | *Syn* |
| MT-ND5 | A13800C | T489P |
| MT-ND5 | G13803A | A490T |
| MT-ND5 | G13803C | A490P |
| MT-ND5 | G13803T | A490S |
| MT-ND5 | C13806A | L491I |
| MT-ND5 | C13808A | *Syn* |
| MT-ND5 | A13815T | T494S |
| MT-ND5 | C13816T | T494I |
| MT-ND5 | C13820A | F495L |
| MT-ND5 | G13825T | G497V |
| MT-ND5 | C13827T | L498F |
| MT-ND5 | T13829C | *Syn* |
| MT-ND5 | C13838A | *Syn* |
| MT-ND5 | G13842T | D503Y |
| MT-ND5 | C13844A | D503E |
| MT-ND5 | C13850A | N505K |
| MT-ND5 | C13850T | *Syn* |
| MT-ND5 | T13855C | L507P |
| MT-ND5 | C13859A | *Syn* |
| MT-ND5 | A13860C | N509H |
| MT-ND5 | A13870C | K512T |
| MT-ND5 | C13880A | *Syn* |
| MT-ND5 | T13885A | L517Q |
| MT-ND5 | C13889T | *Syn* |
| MT-ND5 | C13901T | *Syn* |
| MT-ND5 | C13903A | S523Y |
| MT-ND5 | C13913A | *Syn* |
| MT-ND5 | A13921C | Y529S |
| MT-ND5 | C13923A | P530T |

| | | |
|---|---|---|
| MT-ND5 | T13925C | *Syn* |
| MT-ND5 | T13930C | I532T |
| MT-ND5 | C13931A | I532M |
| MT-ND5 | C13935A | H534N |
| MT-ND5 | A13936C | H534P |
| MT-ND5 | C13940A | *Syn* |
| MT-ND5 | C13940T | *Syn* |
| MT-ND5 | C13940G | *Syn* |
| MT-ND5 | T13945C | I537T |
| MT-ND5 | C13946A | I537M |
| MT-ND5 | C13946T | *Syn* |
| MT-ND5 | A13951G | Y539C |
| MT-ND5 | G13956C | G541R |
| MT-ND5 | G13956T | G541C |
| MT-ND5 | C13958A | *Syn* |
| MT-ND5 | C13966A | T544K |
| MT-ND5 | C13966T | T544M |
| MT-ND5 | G13967T | *Syn* |
| MT-ND5 | C13977A | L548M |
| MT-ND5 | C13981A | P549H |
| MT-ND5 | C13981T | P549L |
| MT-ND5 | C13982A | *Syn* |
| MT-ND5 | C13992T | *Syn* |
| MT-ND5 | C13998A | L555M |
| MT-ND5 | C14002A | T556N |
| MT-ND5 | C14003A | *Syn* |
| MT-ND5 | G14005T | W557L |
| MT-ND5 | C14035A | S567Ter |
| MT-ND5 | C14035T | S567L |
| MT-ND5 | C14035G | S567W |
| MT-ND5 | C14043A | Q570K |
| MT-ND5 | T14047C | I571T |
| MT-ND5 | C14053A | T573N |
| MT-ND5 | C14053T | T573I |
| MT-ND5 | C14056A | S574Y |
| MT-ND5 | C14057A | *Syn* |
| MT-ND5 | C14071A | T579N |
| MT-ND5 | A14075G | *Syn* |
| MT-ND5 | G14079A | G582S |
| MT-ND5 | G14079C | G582R |
| MT-ND5 | C14081A | *Syn* |
| MT-ND5 | C14091T | L586F |
| MT-ND5 | C14100A | L589I |
| MT-ND5 | C14102A | *Syn* |
| MT-ND5 | T14112C | F593L |
| MT-ND5 | A14117C | *Syn* |
| MT-ND5 | C14118T | L595F |
| MT-ND5 | T14119A | L595H |
| MT-ND5 | C14123A | I596M |
| MT-ND5 | C14129A | *Syn* |
| MT-ND5 | T14140A | I602N |
| MT-ND5 | C14143A | T603K |
| MT-ND6 | T14153C | N174D |
| MT-ND6 | C14156A | G173W |
| MT-ND6 | C14168A | E169Ter |

| | | |
|---|---|---|
| MT-ND6 | C14168T | E169K |
| MT-ND6 | C14174A | V167L |
| MT-ND6 | C14183A | V164L |
| MT-ND6 | C14183T | V164M |
| MT-ND6 | A14184C | *Syn* |
| MT-ND6 | C14186A | G163C |
| MT-ND6 | C14186G | G163R |
| MT-ND6 | G14197T | T159K |
| MT-ND6 | C14221A | W151L |
| MT-ND6 | C14221T | W151Ter |
| MT-ND6 | C14226A | *Syn* |
| MT-ND6 | C14227T | G149E |
| MT-ND6 | C14238A | L145F |
| MT-ND6 | A14239G | L145S |
| MT-ND6 | G14242A | A144V |
| MT-ND6 | G14242T | A144D |
| MT-ND6 | C14243A | A144S |
| MT-ND6 | C14247A | *Syn* |
| MT-ND6 | C14249A | A142S |
| MT-ND6 | G14257T | P139H |
| MT-ND6 | C14265A | *Syn* |
| MT-ND6 | C14265T | *Syn* |
| MT-ND6 | G14267A | R136W |
| MT-ND6 | G14267T | R136Ter |
| MT-ND6 | C14305A | S123I |
| MT-ND6 | C14308A | G122V |
| MT-ND6 | C14312A | V121L |
| MT-ND6 | C14314T | S120N |
| MT-ND6 | C14314G | S120T |
| MT-ND6 | C14328A | *Syn* |
| MT-ND6 | C14328T | *Syn* |
| MT-ND6 | C14342A | G111W |
| MT-ND6 | C14342T | G111Ter |
| MT-ND6 | C14342G | G111R |
| MT-ND6 | C14345A | D110Y |
| MT-ND6 | C14351A | E108Ter |
| MT-ND6 | C14358A | W105C |
| MT-ND6 | C14364A | *Syn* |
| MT-ND6 | C14366A | V103L |
| MT-ND6 | C14366T | V103M |
| MT-ND6 | A14374C | V100G |
| MT-ND6 | C14376A | E99D |
| MT-ND6 | C14379A | M98I |
| MT-ND6 | C14379T | *Syn* |
| MT-ND6 | C14382A | *Syn* |
| MT-ND6 | C14382G | *Syn* |
| MT-ND6 | C14389A | G95V |
| MT-ND6 | C14391A | *Syn* |
| MT-ND6 | A14392C | V94G |
| MT-ND6 | C14393T | V94M |
| MT-ND6 | C14399G | V92L |
| MT-ND6 | A14404T | V90E |
| MT-ND6 | C14412T | *Syn* |
| MT-ND6 | C14420A | G85W |
| MT-ND6 | C14420T | G85Ter |

| | | |
|---|---|---|
| MT-ND6 | C14424A | *Syn* |
| MT-ND6 | C14427A | W82C |
| MT-ND6 | T14430A | *Syn* |
| MT-ND6 | G14431A | A81V |
| MT-ND6 | C14433A | E80D |
| MT-ND6 | C14433T | *Syn* |
| MT-ND6 | T14434C | E80G |
| MT-ND6 | G14437A | P79L |
| MT-ND6 | C14442A | E77D |
| MT-ND6 | C14442T | *Syn* |
| MT-ND6 | C14445A | E76D |
| MT-ND6 | A14449T | I75N |
| MT-ND6 | A14455G | M73T |
| MT-ND6 | G14461A | T71M |
| MT-ND6 | G14461T | T71K |
| MT-ND6 | C14484A | M63I |
| MT-ND6 | T14513A | M54L |
| MT-ND6 | C14518A | G52V |
| MT-ND6 | C14519A | G52C |
| MT-ND6 | T14524A | Y50F |
| MT-ND6 | T14539A | N45I |
| MT-ND6 | T14539C | N45S |
| MT-ND6 | C14541A | *Syn* |
| MT-ND6 | C14541G | *Syn* |
| MT-ND6 | C14556A | *Syn* |
| MT-ND6 | C14558A | G39W |
| MT-ND6 | C14561A | V38F |
| MT-ND6 | C14564A | V37L |
| MT-ND6 | C14567A | G36C |
| MT-ND6 | G14568T | S35Ter |
| MT-ND6 | C14577A | L32F |
| MT-ND6 | C14589A | *Syn* |
| MT-ND6 | G14602A | S24F |
| MT-ND6 | C14607A | K22N |
| MT-ND6 | G14611A | S21F |
| MT-ND6 | G14611T | S21Y |
| MT-ND6 | G14614A | S20F |
| MT-ND6 | C14620A | G18V |
| MT-ND6 | C14621A | G18W |
| MT-ND6 | C14624A | V17L |
| MT-ND6 | C14631A | M14I |
| MT-ND6 | C14631T | *Syn* |
| MT-ND6 | C14636A | V13L |
| MT-ND6 | C14642A | G11C |
| MT-ND6 | C14663A | A4S |
| MT-ND6 | C14663T | A4T |
| MT-ND6 | A14668T | M2K |
| MT-ND6 | C14670A | M1I |
| MT-CYB | C14752A | P3T |
| MT-CYB | C14760A | *Syn* |
| MT-CYB | C14760G | *Syn* |
| MT-CYB | A14768C | N8T |
| MT-CYB | A14768G | N8S |
| MT-CYB | C14770A | P9T |
| MT-CYB | C14793A | H16Q |

| | | |
|---|---|---|
| MT-CYB | C14793T | *Syn* |
| MT-CYB | A14796T | *Syn* |
| MT-CYB | C14819A | S25Y |
| MT-CYB | C14831A | A29E |
| MT-CYB | G14834T | W30L |
| MT-CYB | A14850T | *Syn* |
| MT-CYB | C14853A | *Syn* |
| MT-CYB | G14858A | G38D |
| MT-CYB | G14858T | G38V |
| MT-CYB | G14860A | A39T |
| MT-CYB | G14860T | A39S |
| MT-CYB | G14864A | C40Y |
| MT-CYB | C14865A | C40W |
| MT-CYB | C14866A | L41M |
| MT-CYB | C14871A | I42M |
| MT-CYB | C14872A | L43I |
| MT-CYB | C14872T | L43F |
| MT-CYB | C14875A | Q44K |
| MT-CYB | T14879C | I45T |
| MT-CYB | C14882A | T46N |
| MT-CYB | C14882T | T46I |
| MT-CYB | C14883A | *Syn* |
| MT-CYB | G14888A | G48E |
| MT-CYB | G14888T | G48V |
| MT-CYB | C14896A | L51M |
| MT-CYB | A14909G | Y55C |
| MT-CYB | A14918G | D58G |
| MT-CYB | C14921A | A59D |
| MT-CYB | C14921T | A59V |
| MT-CYB | C14924A | S60Ter |
| MT-CYB | C14947A | H68N |
| MT-CYB | C14949T | *Syn* |
| MT-CYB | T14968A | Y75N |
| MT-CYB | T14970A | Y75Ter |
| MT-CYB | G14975T | W77L |
| MT-CYB | A14977G | I78V |
| MT-CYB | C14982A | I79M |
| MT-CYB | C14982G | I79M |
| MT-CYB | C14985A | *Syn* |
| MT-CYB | C14985T | *Syn* |
| MT-CYB | C14988A | Y81Ter |
| MT-CYB | C14988T | *Syn* |
| MT-CYB | C14989A | L82I |
| MT-CYB | C14989T | L82F |
| MT-CYB | C14989G | L82V |
| MT-CYB | C14992A | H83N |
| MT-CYB | G14995C | A84P |
| MT-CYB | G14995T | A84S |
| MT-CYB | G15004A | A87T |
| MT-CYB | G15004C | A87P |
| MT-CYB | G15004T | A87S |
| MT-CYB | C15008A | S88Ter |
| MT-CYB | T15020A | I92N |
| MT-CYB | C15021A | I92M |
| MT-CYB | G15023T | C93F |

| | | |
|---|---|---|
| MT-CYB | C15024A | C93W |
| MT-CYB | C15024T | *Syn* |
| MT-CYB | C15024G | C93W |
| MT-CYB | C15027A | *Syn* |
| MT-CYB | C15030A | F95L |
| MT-CYB | C15031A | L96M |
| MT-CYB | G15041T | G99V |
| MT-CYB | G15047T | G101V |
| MT-CYB | C15074A | S110Ter |
| MT-CYB | A15079C | T112P |
| MT-CYB | A15079G | T112A |
| MT-CYB | G15083A | W113Ter |
| MT-CYB | G15083T | W113L |
| MT-CYB | C15093A | *Syn* |
| MT-CYB | C15093T | *Syn* |
| MT-CYB | C15102A | *Syn* |
| MT-CYB | C15103A | L120M |
| MT-CYB | C15110A | A122E |
| MT-CYB | C15122A | T126K |
| MT-CYB | C15125A | A127D |
| MT-CYB | C15126A | *Syn* |
| MT-CYB | C15126T | *Syn* |
| MT-CYB | T15127C | F128L |
| MT-CYB | G15133T | G130C |
| MT-CYB | G15134T | G130V |
| MT-CYB | T15136C | Y131H |
| MT-CYB | C15141A | *Syn* |
| MT-CYB | C15142A | L133I |
| MT-CYB | C15154A | Q137K |
| MT-CYB | C15154T | Q137Ter |
| MT-CYB | A15157T | M138L |
| MT-CYB | C15161A | S139Ter |
| MT-CYB | G15167A | W141Ter |
| MT-CYB | G15167T | W141L |
| MT-CYB | C15173T | A143V |
| MT-CYB | C15174A | *Syn* |
| MT-CYB | C15174T | *Syn* |
| MT-CYB | C15185A | T147K |
| MT-CYB | A15188G | N148S |
| MT-CYB | T15196C | S151P |
| MT-CYB | C15200A | A152D |
| MT-CYB | C15204A | I153M |
| MT-CYB | C15204T | *Syn* |
| MT-CYB | A15211T | I156F |
| MT-CYB | G15214A | G157Ter |
| MT-CYB | G15214T | G157W |
| MT-CYB | G15215T | G157V |
| MT-CYB | C15222A | D159E |
| MT-CYB | G15226A | V161I |
| MT-CYB | T15228C | *Syn* |
| MT-CYB | C15229A | Q162K |
| MT-CYB | C15237A | I164M |
| MT-CYB | G15239T | W165L |
| MT-CYB | G15244A | G167S |
| MT-CYB | G15244T | G167C |

| | | |
|---|---|---|
| MT-CYB | C15246A | *Syn* |
| MT-CYB | G15253A | V170M |
| MT-CYB | C15262A | P173T |
| MT-CYB | C15263A | P173H |
| MT-CYB | C15264A | *Syn* |
| MT-CYB | C15264T | *Syn* |
| MT-CYB | A15265C | T174P |
| MT-CYB | C15267A | *Syn* |
| MT-CYB | T15269A | L175H |
| MT-CYB | T15269C | L175P |
| MT-CYB | C15274A | R177Ter |
| MT-CYB | C15285G | *Syn* |
| MT-CYB | C15291T | *Syn* |
| MT-CYB | C15297A | I184M |
| MT-CYB | G15300A | *Syn* |
| MT-CYB | C15302A | P186H |
| MT-CYB | C15303A | *Syn* |
| MT-CYB | T15305C | F187S |
| MT-CYB | C15317A | A191D |
| MT-CYB | C15318A | *Syn* |
| MT-CYB | C15334A | L197I |
| MT-CYB | C15336T | *Syn* |
| MT-CYB | C15337T | *Syn* |
| MT-CYB | T15341C | F199S |
| MT-CYB | G15345C | L200F |
| MT-CYB | G15345T | L200F |
| MT-CYB | C15346A | H201N |
| MT-CYB | C15348A | H201Q |
| MT-CYB | G15349T | E202Ter |
| MT-CYB | G15354C | *Syn* |
| MT-CYB | G15354T | *Syn* |
| MT-CYB | G15355A | G204Ter |
| MT-CYB | G15355T | G204W |
| MT-CYB | G15356T | G204V |
| MT-CYB | C15367A | P208T |
| MT-CYB | C15369A | *Syn* |
| MT-CYB | C15370A | L209M |
| MT-CYB | G15374A | G210E |
| MT-CYB | C15378A | I211M |
| MT-CYB | C15378T | *Syn* |
| MT-CYB | C15381A | *Syn* |
| MT-CYB | C15384A | *Syn* |
| MT-CYB | T15398C | I218T |
| MT-CYB | C15410A | P222H |
| MT-CYB | C15410T | P222L |
| MT-CYB | T15422A | I226N |
| MT-CYB | C15423A | I226M |
| MT-CYB | G15427A | D228N |
| MT-CYB | C15432A | *Syn* |
| MT-CYB | C15435A | *Syn* |
| MT-CYB | C15435T | *Syn* |
| MT-CYB | C15438A | *Syn* |
| MT-CYB | C15450A | F235L |
| MT-CYB | T15455A | L237H |
| MT-CYB | C15456G | *Syn* |

| | | |
|---|---|---|
| MT-CYB | G15465C | M240I |
| MT-CYB | G15465T | M240I |
| MT-CYB | C15489T | *Syn* |
| MT-CYB | C15490A | L249I |
| MT-CYB | C15490T | L249F |
| MT-CYB | C15492A | *Syn* |
| MT-CYB | C15493A | L250M |
| MT-CYB | C15501A | D252E |
| MT-CYB | C15501T | *Syn* |
| MT-CYB | A15514C | T257P |
| MT-CYB | C15516A | *Syn* |
| MT-CYB | C15516T | *Syn* |
| MT-CYB | C15521A | A259D |
| MT-CYB | C15521T | A259V |
| MT-CYB | C15526A | P261T |
| MT-CYB | C15541A | P266T |
| MT-CYB | C15542A | P266H |
| MT-CYB | C15542T | P266L |
| MT-CYB | A15551T | K269M |
| MT-CYB | G15556T | E271Ter |
| MT-CYB | G15560A | W272Ter |
| MT-CYB | G15560T | W272L |
| MT-CYB | T15585A | I280M |
| MT-CYB | T15587A | L281H |
| MT-CYB | T15587C | L281P |
| MT-CYB | C15597A | *Syn* |
| MT-CYB | G15613A | G290S |
| MT-CYB | G15613T | G290C |
| MT-CYB | G15614A | G290D |
| MT-CYB | G15614T | G290V |
| MT-CYB | C15619A | L292I |
| MT-CYB | C15619T | L292F |
| MT-CYB | C15625A | L294M |
| MT-CYB | C15625T | *Syn* |
| MT-CYB | C15635A | S297Y |
| MT-CYB | C15636A | *Syn* |
| MT-CYB | C15645A | I300M |
| MT-CYB | C15650A | A302E |
| MT-CYB | C15657A | I304M |
| MT-CYB | C15658A | P305T |
| MT-CYB | C15660G | *Syn* |
| MT-CYB | C15663A | I306M |
| MT-CYB | C15667A | H308N |
| MT-CYB | C15667T | H308Y |
| MT-CYB | C15700A | P319T |
| MT-CYB | C15709A | Q322K |
| MT-CYB | C15713A | S323Ter |
| MT-CYB | C15724A | L327I |
| MT-CYB | C15724T | L327F |
| MT-CYB | C15731A | A329D |
| MT-CYB | C15731T | A329V |
| MT-CYB | C15734A | A330E |
| MT-CYB | T15740C | L332P |
| MT-CYB | C15748A | L335M |
| MT-CYB | C15753T | *Syn* |

| | | |
|---|---|---|
| MT-CYB | C15769A | P342T |
| MT-CYB | C15777A | S344Ter |
| MT-CYB | T15784C | F347L |
| MT-CYB | C15792A | I349M |
| MT-CYB | C15799A | Q352K |
| MT-CYB | G15805C | A354P |
| MT-CYB | G15805T | A354S |
| MT-CYB | C15809A | S355Y |
| MT-CYB | C15831A | I362M |
| MT-CYB | C15832A | L363M |
| MT-CYB | C15832T | *Syn* |
| MT-CYB | C15838T | *Syn* |
| MT-CYB | C15844A | P367T |
| MT-CYB | C15845A | P367Q |
| MT-CYB | C15854A | S370Y |
| MT-CYB | C15856A | L371M |
| MT-CYB | C15856G | L371V |
| MT-CYB | T15857C | L371P |
| MT-CYB | T15880A | W379Ter |
| MT-CYB | G15883T | A380S |
| MT-CYB | C15884A | A380D |
| MT-CYB | C15885A | *Syn* |

f human breast normal cells determined via Duplex Sequencing.
